# Identification of a Central Neural Circuit that Regulates Anxiety-induced Bone Loss

**DOI:** 10.1101/812875

**Authors:** Fan Yang, Yunhui Liu, Shanping Chen, Zhongquan Dai, Dazhi Yang, Dashuang Gao, Jie Shao, Yuyao Wang, Ting Wang, Zhijian Zhang, Lu Zhang, William W Lu, Jie Tu, Yinghui Li, Liping Wang

## Abstract

The homeostasis of bone metabolism is finely regulated by the central nervous system and recent studies have suggested that mood disorders, such as anxiety, are closely related to bone metabolic abnormalities; however, our understanding of central neural circuits regulating bone metabolism is still largely limited. In this study, we first demonstrate that confined isolation of human participants under normal gravity resulted in decreased bone density and elevated anxiety levels. We then used an established mouse model to dissect the neural circuitry regulating anxiety-induced bone loss. Combining electrophysiological, optogenetic and chemogenetic approaches, we demonstrate that GABAergic neural circuitry in ventromedial hypothalamus (VMH) modulates anxiety-induced bone loss; importantly, the GABAergic input in VMHdm arose from a specific group of somatostatin neurons in the bed nucleus of the stria terminalis (BNST), which is both indispensable for anxiety-induced bone loss and able to trigger bone loss in the absence of stressors. VGLUT2 neurons in Nucleus tractus solitaries (NTS) and peripheral sympathetic system were employed by this BNST-VMH neural circuit to regulate anxiety-induced bone loss. Overall, we uncovered new GABAergic neural circuitry from the forebrain to hypothalamus, used in the regulation of anxiety-induced bone loss, and revealed a population of somatostatin neurons in BNST not previously implicated in bone mass regulation. These findings thus identify the underlying central neural mechanism of psychiatric disorders, such as anxiety, that influences bone metabolism at the circuit level.

**One Sentence Summary:** Identification of a new GABAergic neural circuit from forebrain to hypothalamus used for regulation of anxiety-induced bone loss.

## Introduction

Mood disorders, such as depression and anxiety, are closely related with bone metabolism abnormalities (*1–3*) and can lead to low bone mass and increased risk of fracture (*4, 5*). Another well-established cause of bone loss is microgravity during spaceflight (*6*); however, decreased bone formations in astronauts occurred (*7, 8*) or progressively deteriorated (*9*) under normal gravity following space missions. Chronic stressors in extreme microenvironments may not only induce psychological changes in crew but also continuously disturb peripheral metabolism through the central nervous system (*10, 11*). Based on these findings, we reasoned that stress-induced psychological disorders such as anxiety might directly contribute to bone loss through the central nervous system; however, the exact neural mechanism behind the anxiety-induced bone loss is not clear.

Bone metabolism is a dynamic physiological process that is finely regulated by the brain (*12*); central signals from the brain exert important influences on bone metabolism (*13–15*). For example, leptin signaling in the hypothalamus inhibits bone formation through the sympathetic nervous system (*14*) and orexin, which is produced in the hypothalamus, is a critical regulator of skeletal homeostasis and exerts regulation of bone mass (*16*). The hypothalamus is an important neural control center relaying information from higher centers in the forebrain to regulate body metabolism (*17*). Among many hypothalamic nuclei, the ventromedial hypothalamus (VMH) is a distinct region that is important in regulating emotion (*18, 19*), energy balance (*13, 20*) and bone metabolism (*15, 21*). Knocking-out SF1 neurons, which are present in the VMH, results in significantly increased anxiety-like behaviors (*22*), and optogenetic activation of dorsal VMH promotes a variety of context-dependent defense-like and autonomic responses (*23*). Moreover, inactivation of CaMKKβ in SF1-expressing neurons in the VMH results a severe low bone phenotype (*21*), whilst serotoninergic projections from the raphe nucleus to the VMH may regulate SF1 neurons via Htr2c receptors to promote bone accrual (*15*). Manipulations of the transcription factor, AP1, in VMH SF1-expressing neurons leads to an increase in energy and a decrease in bone density, suggesting that the VMH regulates bone homeostasis and energy metabolism through disparate neural pathways (*24*). Collectively, these findings suggest that differential neural pathways direct VMH neurons to regulate emotions and bone metabolism in a top-down manner. However, how the specific neural circuit involving the VMH that integrates anxiety information and regulates downstream bone metabolism is unknown.

In this study we found, firstly, that chronic stress in crewmembers simulating a long-term space mission resulted in both decreased bone density and elevated anxiety levels under normal gravity. We then used an established mouse model to determine the neural circuitry underlying anxiety-induced bone loss. Secondly, by combining electrophysiological, optogenetic and chemogenetic approaches, we found that the GABAergic neural circuity in VMH is responsive in the anxiety-induced bone loss; importantly, the GABAergic input in VMHdm originate in a specific group of somatostatin neurons in the posterior region of the bed nucleus of the stria terminalis (BNST), which is both indispensable for and able to drive the anxiety-induced bone loss. The Nucleus tractus solitaries (NTS) and peripheral sympathetic system were employed by the BNST-VMH neural circuitry to regulate the anxiety-induced bone loss. Our study thus identified GABAergic neural circuitry from the forebrain to the hypothalamus used in the regulation of anxiety-induced bone loss, revealed a population of somatostatin neurons in BNST not previously implicated in bone mass regulation, and provides new insights in understanding how anxiety disorders can influence bone metabolism from a neural circuits level.

## Results

### 180 days of chronic stress in crewmembers results in decreased bone density and elevated anxiety level

To investigate the relationship between anxiety and bone loss, we first performed a Controlled Ecological Life Support System (CELSS) integrated experiment where four crewmembers spent 180 days in an isolated habitat mimicking a space station under normal gravity. Weight, anxiety level and biochemical parameters were measured in all participants every 30 days during the mission (Figure 1A). While weight and cortisol levels were stable for each of the four crewmembers during the mission (Figure 1B, C), the Self-Rating Anxiety Scale (SAS) measurements showed increased scores during the prolonged stay in the confined space, which were statistically significant (Figure 1D). There was also a significant increase in epinephrine and norepinephrine (NE) levels during the 180 days (Figure 1E, F), reflecting the elevated level of reported crewmember anxiety. Bone density analysis revealed a statistically significant decrease in average bone mineral density (BMD) in femur, femur neck and the lumbar vertebrae in all participants during the mission (Table 1, Figure 1I-K). Bone formation markers alkaline phosphatase (ALP) and Procollagen I carboxy-terminal propeptide (PICP) also continuously decreased in all participants (Figure 1G, H). These data consistently show that the 180 days of chronic stress on four crewmembers resulted in elevated anxiety levels and decreased bone density.

**Fig. 1.**
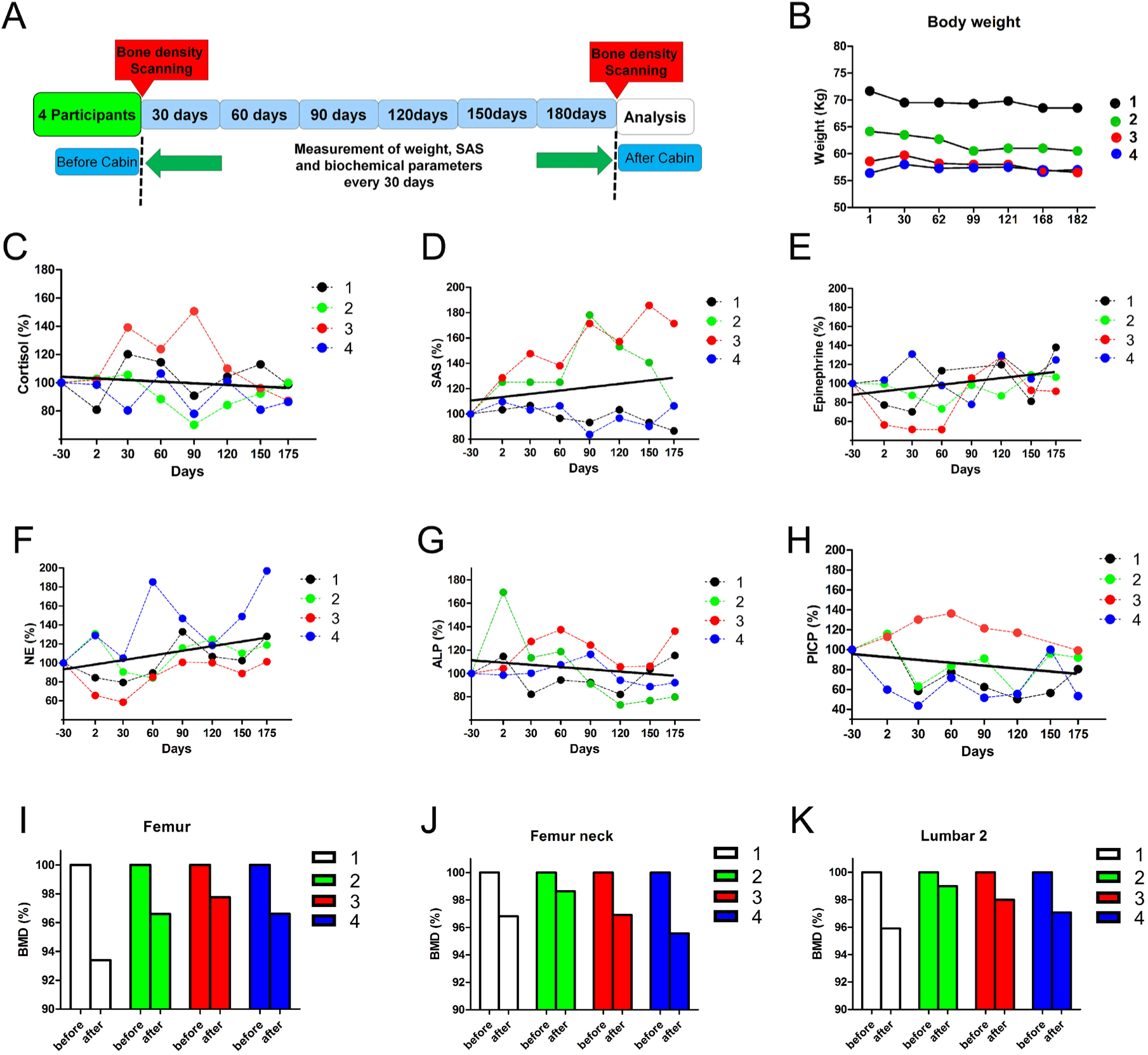
Chronic stress in four crewmembers resulted in decreased bone density and elevated anxiety levels. **(A)** Schematic of the experimental protocol used in the 180-day Controlled Ecological Life Support System integrated experiment. Biochemical parameters were measured every 30 days for all crewmembers; bone mineral density (BMD) was measured before and after the experiment. **(B)** During the mission, normal bodyweight was maintained in all 4 crewmembers. **(C)** Cortisol levels at 30-day intervals during the mission (n=4). **(D)** Self-Rating Anxiety Scale (SAS) scores every 30 days during the prolonged stay in the confined space (n=4). **(E)** Epinephrine levels every 30 days during the confined isolation (n=4). **(F)** Norepinephrine (NE) levels every 30 days during the confined isolation (n=4). **(G)** Bone formation marker alkaline phosphatase (ALP) levels every 30 days during the confined isolation (n=4). **(H)** Procollagen I carboxy-terminal propeptide (PICP) levels at 30-day intervals during the mission (n=4). **(I)** BMD in femur of the 4 crewmembers before and after confinement. **(J)** BMD in femur neck of the 4 participants before and after confinement. **(K)** BMD in the 2nd lumbar vertebrae of the 4 participants before and after the confinement.

**Table 1.** Changes in bone mineral density (BMD) in different anatomical regions in the four crewmembers after the experiment.

|  | BMD Changes Rate (A-B)/B*100% |  |  |  |  |  |
| --- | --- | --- | --- | --- | --- | --- |
| **Participants** | **1** | **2** | **3** | **4** | **Average** | **SD** |
| **Lumbar 1** | -0.25% | 4.53% | -4.06% | 1.02% | **0.31%** | 3.55% |
| **Lumbar 2** | -4.10% | -1.01% | -2.01% | -2.92% | **-2.51%** | 1.32% |
| **Lumbar 3** | -5.64% | 0.34% | 1.80% | -1.87% | **-1.34%** | 3.24% |
| **Lumbar 4** | -1.82% | -2.59% | 2.96% | 0.08% | **-0.34%** | 2.47% |
| **L1-L2** | -2.29% | 1.63% | -2.84% | -0.98% | **-1.12%** | 1.99% |
| **L1-L3** | -3.56% | 1.20% | -1.18% | -1.36% | **-1.23%** | 1.95% |
| **L1-L4** | -3.13% | 0.00% | 0.08% | -0.98% | **-1.01%** | 1.49% |
| **L2-L3** | -4.89% | -0.27% | 0.00% | -2.44% | **-1.9%** | 2.27% |
| **L2-L4** | -3.77% | -1.14% | 1.13% | -1.59% | **-1.34%** | 2.01% |
| **L3-L4** | -3.63% | -1.20% | 2.47% | -0.85% | **-0.80%** | 2.51% |
| **Femur neck** | -11.23% | -1.37% | -3.10% | -4.43% | **-5.03%** | 4.32% |
| **Femur** | -6.61% | -3.40% | -2.24% | -3.39% | **-3.91%** | 1.88% |
| **Radius** | -0.27% | -2.17% | 0.28% | -2.56% | **-1.18%** | 1.39% |
| **Unla** | 1.80% | -4.29% | 0.38% | -0.77% | **-0.72%** | 2.60% |
| BMD: Bone Mineral Density; B: before cabin; A: after cabin |  |  |  |  |  |  |

### GABAergic neural circuitry in dorsomedial VMH is actively involved in a mouse model of anxiety-induced bone loss

To dissect the neural mechanism of anxiety induced bone loss, we exposed mice to chronic mild stressors mimicking the CELSS and thus established a mouse model of chronic stress-induced anxiety (Figure 2A). After eight weeks of exposure, we assessed anxiety-related behavior using an open field (OF) test and an elevated plus maze (EPM) test. In the OF test, mice in the stress group exhibited significantly less entries in the central area and spent less time exploring the central area compared with the control group (Figure 2B). In the EPM test, mice in the stress group exhibited significantly less entries and spent less time in the open arms compared with the control group (Figure 2C). Micro-CT analysis of the trabecular bones in the proximal tibia revealed that mice in the stress group displayed an obvious low bone mass phenotype (Figure 2D). The trabecular bone volume/tissue volume (BV/TV) ratio in the stress group was 18% lower than the control group. The trabecular number (TbN) was 15% lower, accompanied by 13% higher trabecular separation (Tb.Sp) in the stress group compared with control group (Figure 2E), illustrating decreased bone mass in the stress group. Linear regression analysis using BV/TV and time spent in central area/open arms revealed a strong relationship and inverse association between BV/TV and parameters reflecting level of anxiety (Figure 2F). No statistical differences in total distance traveled in OF, serum cortisol levels or body weight were observed between the stress and control groups (Figure S1B-S1D). However, NE levels were higher and ALP levels were lower in the stress group compared with the control group, and these results were statistically significant (Figure S1E and S1F).

**Fig. 2.**
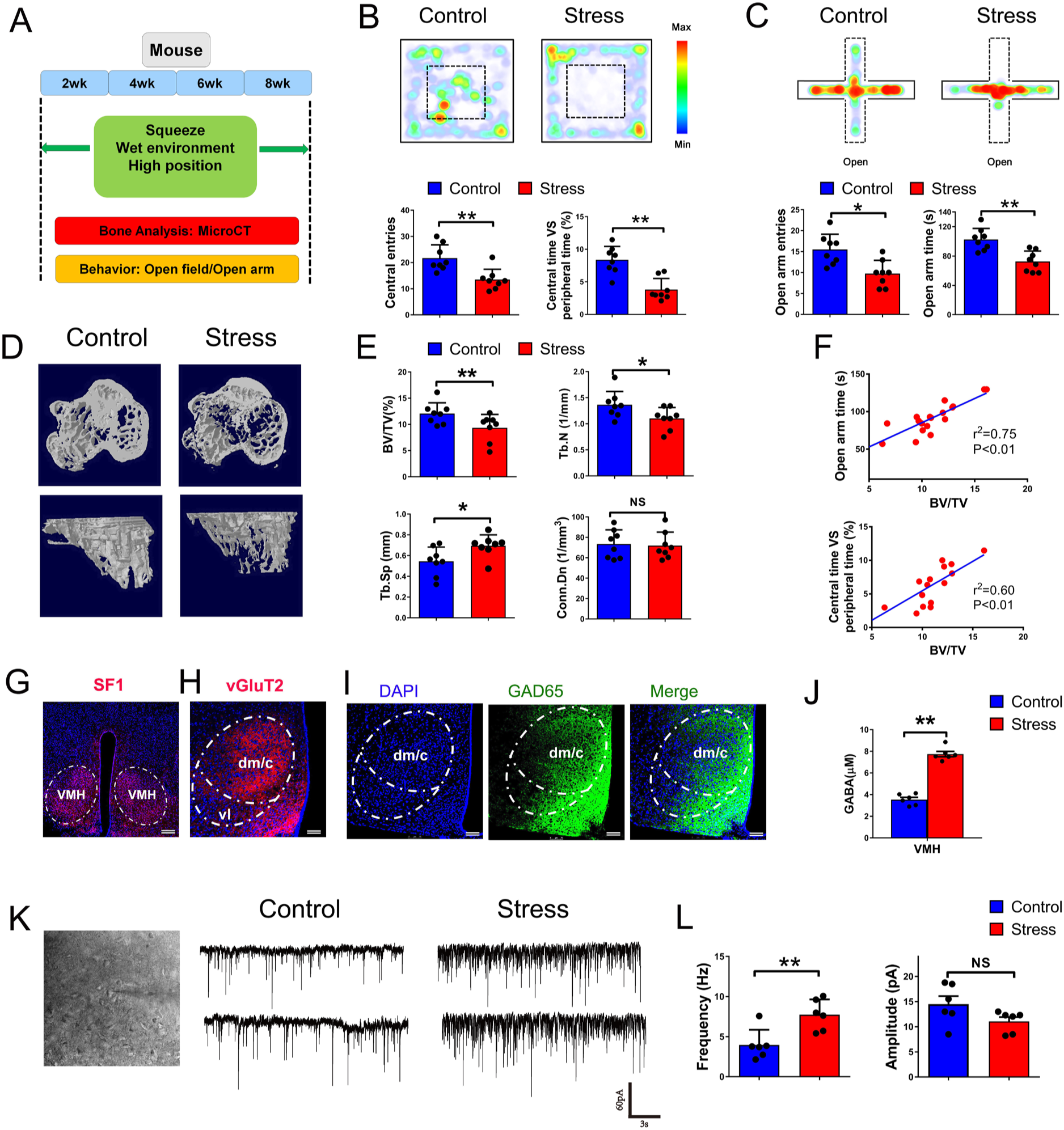
A mouse model of chronic stress induced anxiety-related behavior and bone loss. **(A)** Schematic for establishing the mouse model; C57BL/6 mice were exposed to chronic stressors for 8 weeks, then behavioral tests and bone analysis were conducted at the end of the 8 weeks. **(B)** Open field (OF) test comparing the stress and control groups; mice in the stress group exhibited significantly fewer entries in central zone and spent less time exploring the central area. Values represent mean±SEM (n=8 per group; \*\**p*<0.01; Student’s t test). **(C)** Elevated plus maze (EPM) test comparing stress and control groups; mice in the stress group exhibited significantly fewer entries and spent less time in the open arms. Values represent mean±SEM (n=8 per group; * *p*<0.05; \*\**p*<0.01; Student’s t test). **(D)** Representative coronal (top) and sagittal (bottom) plane images of proximal tibia; mice in the stress group displayed a low bone-mass phenotype. **(E)** MicroCT analysis of the parameters including trabecular bone volume/tissue volume (BV/TV), the trabecular number (TbN), trabecular separation (Tb.Sp), and connectivity density (Conn. Dn) in the stress and control groups. Values represent mean±SEM (n=8 per group; * *p*<0.05; \*\**p*<0.01; NS, not significant; Student’s t test). **(F)** Correlation analysis using BV/TV and time spent in central area/open arm that reflect the level of anxiety (n=16). **(G)** Immunostaining of SF1 neurons in the VMH; scale bar, 150 μm. **(H)** Immunostaining of vesicular glutamate transporter (vGlut2) in dorsomedial region of the VMH (VMHdm); scale bar, 75 μm. **(I)** Immunostaining of GAD65-positive GABAergic projections in the VMHdm; scale bar, 75 μm. **(J)** Quantification of GABA levels in the VMHdm region in stress group and control groups; values represent mean ± SEM (n=6 per group; \*\**p*<0.01; Student’s t test). **(K)** Representative inhibitory postsynaptic currents (IPSCs) of SF1 neurons in the VMHdm in the stress and control groups. **(L)** Quantification of the frequency and amplitude of IPSCs from SF1 neurons in the stress and control groups; values represent mean±SEM (n=6 per group; \*\**p*<0.01; NS, not significant; Student’s t test).

The dorsomedial VMH (VMHdm) is important in regulating anxiety (*23*), energy balance (*13*) and bone metabolism (*25*). The major type of neurons in the VMHdm are SF1 neurons, which are mainly glutamatergic (Figure 2G and 2H) and thus, we next began to dissect the neural mechanism from the VMHdm. Dense GAD65-positive GABAergic projections were observed in the VMHdm (Figure 2I). Microdialysis revealed significantly higher VMHdm GABA levels in the stress group compared with the control group (Figure 2J). We then recorded inhibitory postsynaptic currents (IPSCs) of SF1 neurons in the VMHdm and found a statistically significant elevated frequency of spontaneous IPSCs in the stress group compared with the control group (Figure 2K and 2L); both the frequency and amplitude of IPSCs were completely blocked by 50 μM of bicuculline (Figure S1G-S1I), suggesting that GABAergic neural projections in the VMHdm play an important role in the mouse model of anxiety-induced bone loss.

### Activation of the GABAergic projections in the VMHdm inhibits the firing of SF1 neurons and induces anxiety-like behavior and bone loss

To interrogate the function of GABAergic projections in the VMHdm during anxiety induced bone loss, we used transgenic mice expressing ChR2 under the promoter of the conditional allele of the vesicular GABA transporter (VGAT) (Figure 3A). VGAT signals surrounded SF1 neurons and the distributive pattern of VGAT matched that of GAD65, which stains GABAergic axon terminals (Figure S2A). We recorded IPSCs of SF1 neurons and found that blue light induced excitation of GABAergic terminals, resulting in fast IPSCs recorded from SF1 neurons (Figure 3B). The frequency and amplitude of IPSCs induced by blue light was blocked by 50 μM of bicuculline (Figure S2B and S2C). Patch-clamp recordings combined with single-cell RT-PCR verified that blue light can indeed inhibit the spontaneous firing of SF1 neurons that express the genes *vglut2* and *Sf1* but not *vgat* (Figure S2D and S2E).

**Fig. 3.**
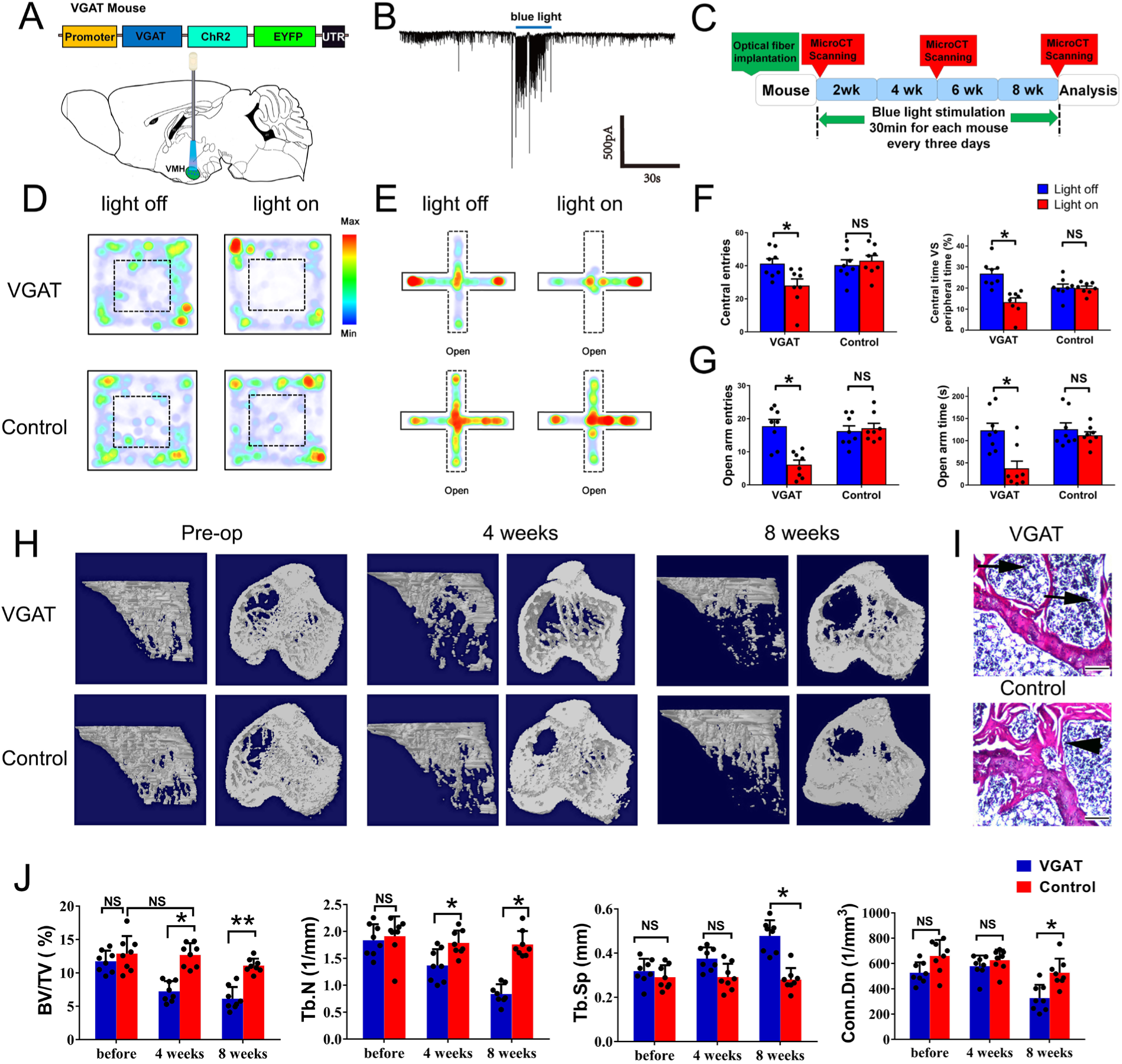
Activation of GABAergic projections in the VMHdm induced anxiety-like behaviors and bone loss. **(A)** Optical fibers were implanted into the VMHdm of VGAT-ChR2 mice and the region was illuminated with blue light. **(B)** Electrophysiological recording of a typical inhibitory postsynaptic current (IPSC) from SF1 neurons during blue light stimulation. **(C)** Schematic showing the schedule of blue light stimulation and bone analysis with intermittent MicroCT scanning. **(D)** Open field test of VGAT mice and control mice before and during blue light stimulation; VGAT mice displayed obvious anxiety-like behavior. **(E)** Elevated plus maze test of VGAT and control mice before and during blue light stimulation; VGAT mice displayed obvious anxiety-like behavior. **(F)** Quantification of the entries to the central area and time spent in the central area before and during blue light stimulation in the VGAT and control groups; values represent mean ± SEM (n=8 per group; * *p*<0.05; NS, not significant; one-way analysis of variance (ANOVA) with Bonferroni correction for multiple comparisons test). **(G)** Quantification of the entries and time spent in the open arms in the VGAT and control groups before and during blue light stimulation. Values represent mean±SEM (n=8 per group; * *p*<0.05; NS, not significant; One-way analysis of variance (ANOVA) with Bonferroni correction for multiple comparisons test). **(H)** MicroCT analysis of VGAT and control group bone structure before, during and after blue light stimulation: a significantly low bone mass phenotype was observed in the VGAT group at 4 and 8 wks after light stimulation began. **(I)** Hematoxylin and eosin (H&E) staining of the proximal tibia in the VGAT and control groups at 8 wks after the stimulation. **(J)** MicroCT analysis of the parameters including trabecular bone volume/tissue volume (BV/TV), the trabecular number (TbN), trabecular separation (Tb.Sp), and connectivity density (Conn. Dn) in the VGAT and control groups. Values represent mean±SEM (n=8 per group; * *p*<0.05; \*\**p*<0.01; NS, not significant; One-way analysis of variance (ANOVA) with Bonferroni correction for multiple comparisons test).

To further investigate the causal role of the activation of GABAergic projections in the VMHdm during anxiety-induced bone loss, we assessed anxiety behavior and evaluated bone structure changes following activation of the VMHdm in VGAT-ChR2 mice (Figure 3C). After a series of light stimulation, GABA levels in the VMHdm were significantly higher in the VGAT group compared with the control group (Figure S2F). Both the OF test and the EPM test showed that VGAT mice displayed obvious anxiety-like behavior compared with the control group during the of the light-on phase (Figures 3D and 3E) and on the OF test there were significantly fewer entries to the central area and less time spent in central area in the VGAT group when the blue light was turned on (Figure 3F). Additionally, the VGAT group also had a significantly fewer entries and time spent on the open arms in the EPM test compared with the control group during the light-on phase (Figure 3G). Interestingly, micro-CT analysis revealed continuous bone loss in VGAT group over the weeks of light stimulation (Figure 3H). Before light stimulation there were no significant differences in BV/TV, TbN, TbSp nor connectivity density (Conn) between the VGAT and control groups. However, four weeks after the light stimulation began, we observed an obvious low bone mass phenotype in the VGAT group (Figure 3J); the VGAT group had a 15% lower BV/TV than the control group and TbN was 18% lower (Figure 3J). Eight weeks after light stimulation began, the VGAT group had an 18% lower BV/TV and a 16% lower TbN than the control group, whereas Tb.Sp was 18% higher (Figure 3J). Histological analysis at eight weeks confirmed the loss of trabecular bone and larger medulla cavity of the proximal tibia in VGAT mice (Figure 3I). Taken together, these data demonstrate that activation of the GABAergic neural terminals in the VMHdm was able to induce the anxiety-behavior and low bone mass phenotype in the absence of additional anxiety, supporting a direct causal role of VMHdm GABAergic projection activation in anxiety-induced bone loss.

### Somatostain neurons send GABAergic projections from the BNST to the VMHdm

To investigate the source of GABAergic input to the VMHdm region, a retrograde virus was used to trace the neuronal connections of SF1 neurons in the VMH region (Figure 4A). Efficient gene delivery was obtained in the VMH region and retrograde mono-synaptic transfer of mCherry allowed the mapping of different brain regions connecting to VMH (Figure 4B). We found that the main upstream inputs to the VMH included hypothalamic regions, such as the paraventricular nucleus (PVN) and the anterior hypothalamic nucleus (AHC) (Figure 4B and 4C). Importantly, we also observed dense signals in both lateral dorsal and posterior regions of the bed nucleus of stria terminalis (BNST), which play a crucial role in stress-related disorders and adaptive anxiety behaviors (Figure 4B). To confirm that the BNST sends GABAergic inputs to the VMHdm, we injected the AAV-Ef1a-DIO-mcherry virus into the BNST region of GAD-Cre mice and observed GABAergic neural projections in the VMHdm region (Figure S3A and S3B). Further confirmation of neuronal activation both in the lateral dorsal and posterior regions of the BNST in our established mouse model came from c-fos staining (Figure 4D) and there was a statistically significant larger number of c-fos positive cells in the anxiety group than the control group (Figure 4E), suggesting that the BNST send GABAergic inputs to the VMHdm to regulate anxiety-induced bone loss. The BNST is comprised of multiple sub-divisions and contains heterogeneous neuronal subpopulations that perform different functions (*26*). To determine the specific neuronal subpopulations projecting to the VMHdm that mediate anxiety-induced bone loss, we started by dissecting the types of GABAergic terminals in the VMHdm region. Compared with parvalbumin (PV) signals and cholecystokinin (CCK) signals, we observed strong somatostatin nerve terminal signals that were making synaptic contact with SF1 neurons in the VMHdm (Figure 4F and 4G). To determine the specific source of somatostatin from the BNST, we injected AAV9 virus expressing mCherry into the VMHdm of SOM-Cre mice and found a specific population of SOM-positive neurons that was positive-labeled in a posterior region (BSTLP) of BNST (Figure S3C). Combining monosynaptic tracing using rabies virus with in-situ hybridization of somatostatin mRNA, we confirmed that the identified SOM neurons in the BSTLP region indeed send projections to innervate SF1 neurons in VMHdm (Figure 4H and 4K). We also observed dense SOM terminals in the VMHdm region when the AAV9 virus was injected into the BSTLP region (Figure 4I, 4J and 4L). These data consistently show a specific population of SOM neurons in the BSTLP region that send GABAergic neural projections to innervate VMHdm SF1 neurons.

**Fig. 4.**
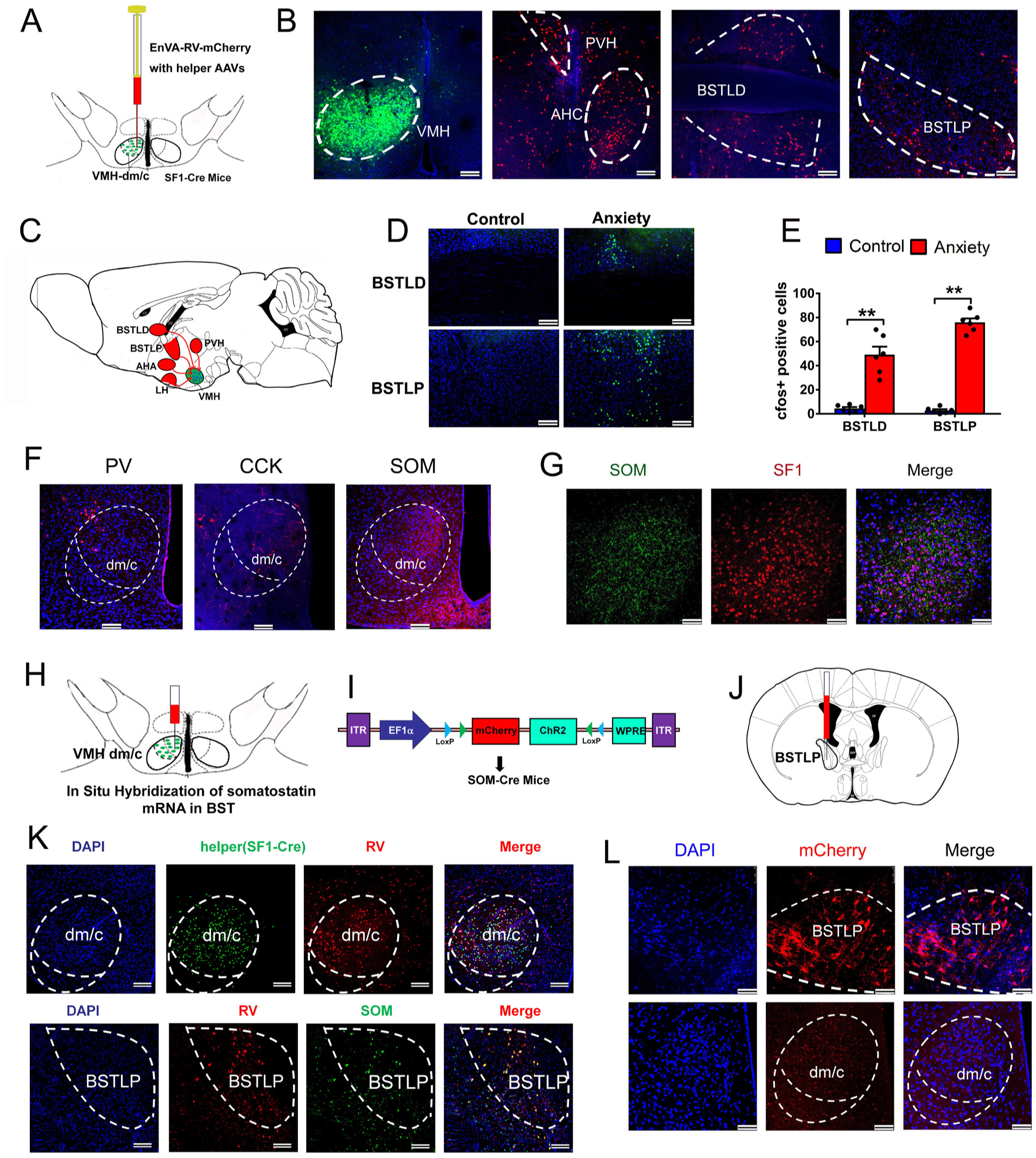
Somatostain neurons in posterior BNST send GABAergic projections to VMHdm region. **(A)** chematic of rabies-based monosynaptic retrograde tracing from SF1 neurons in the VMHdm of SF1-Cre mice; the VMH of SF1-Cre mice was transduced unilaterally with Cre-dependent AAVs encoding TVA-EGFP fusion protein and rabies virus glycoprotein, followed 3 wk later by EnvA-pseudotyped, glycoprotein-deleted mCherry expressing rabies virus. **(B)** Representative images of the VMH region with the helper AAV virus expressing eYFP, and of RV-labeled cells (red) in different brain regions; PVH, paraventricular nucleus; AHC, anterior hypothalamic nucleus; BSTLD, lateral dorsal bed nucleus of stria terminalis; and BSTLP, lateral posterior bed nucleus of stria terminalis; scale bar, 100 μm. **(C)** Schematic mapping of neural circuits from different brain regions to the VMH through monosynaptic neural connection. **(D)** c-fos staining of BSTLD and BSTLP regions in the anxiety and control groups; scale bar, 100 μm. **(E)** Quantification of c-fos positive cells in anxiety and control groups (n=3 mice per group; \*\**p*<0.01; one-way analysis of variance (ANOVA) with Bonferroni correction for multiple comparisons test). **(F)** Immunostaining of neural projections in VMH: PV, parvalbumin; CCK, cholecystokinin; SOM, somatostatin, scale bar, 100 μm. **(G)** Double staining of SF1 neurons and SOM-positive projections in VMHdm; scale bar, 50 μm. **(H)** The rabies virus was injected unilaterally into the dorsomedial region of the VMH in SF1-Cre mice, and in-situ hybridization of somatostatin mRNA in the posterior region of the BNST. **(I)** Schematic of AAV9 expressing mCherry and ChR2 under the EF1α promoter. **(J)** Schematic for the virus injection into the lateral posterior region of BNST of SOM-Cre mice. **(K)** The rabies virus was injected unilaterally into the VMHdm in SF1-Cre mice (up panel); in-situ hybridization of somatostatin mRNA in the BSTLP region (bottom panel); RV signals and somatostatin mRNA were co-localized in the BSTLP region. **(L)** Representative image of SOM positive neurons in the BSTLP (top) and SOM positive neural projections in the VMHdm (bottom) after injection into the BSTLP region; scale bar, 100 μm.

### Activation of SOM neural projections from BNST to VMH induces anxiety-like behavior and bone loss

To determine whether activation of specific SOM inputs from BSTLP was able to cause anxiety-induced bone loss, we injected AAV-Ef1α-DIO-ChR2-mcherry virus into the BSTLP region and used blue light to illuminate the VMHdm region (Figure 5A). GABA levels in the VMHdm were significantly higher in the SOM-ChR2 group compared to controls (Figure 5B), and SOM neurons were activated in the SOM-ChR2 group after light stimulation (Figure 5D). Electrophysiological recordings showed that 20 Hz pulses of blue light inhibited the firing of SF1 neurons in the VMHdm (Figure 5C). We then assessed anxiety-related behavior following light stimulation of SOM neural projections. The mCherry-ChR2 group spent less time and made fewer entries into the center during an OF test compared to the mCherry control group (Figure 5E). In an EPM test, the mCherry-ChR2 group also made less entries to and spent less time in the open arms compared to the control group, which were statistically significant (Figure 5F). Thus, the mCherry-ChR2 group displayed increased anxiety-like behavior following light stimulation. Four weeks after this light stimulation, bone structure was evaluated and mice in mCherry-ChR2 group had a phenotype with a lower bone mass than the mCherry control group, which was statistically significant (Figure 5G): 20% lower BV/TV, 25% lower TbN and 20% higher Tb.Sp (Figure 5H). We found no statistical differences between groups in total distance traveled in the OF test or in body weight (Figure S3D). To confirm the specific function of SOM neural terminals, we optogenetically activated the PV-positive neural terminals and CCK-positive terminals in the VMHdm region using PV-Cre mice and CCK-Cre mice, respectively (Figure S4A and S4B; Figure S5A and S5B). In both PV-Cre and CCK-Cre mice, there was no difference in central area time and central entries in the OF following light stimulation, compared to their respective control groups (Figure S4C and S4D; Figure S5C and S5D) and micro-CT analysis of bone revealed no changes in BV/TV, TbN or TbSp (Figure S4E and S4F; Figure S5E and S5F). Together, our data indicate that activation of the specific SOM inputs from the BNST to the VMHdm was able to induce anxiety and bone loss, implicating the specificity of the population of SOM neurons in regulating anxiety-induced bone loss.

**Fig. 5.**
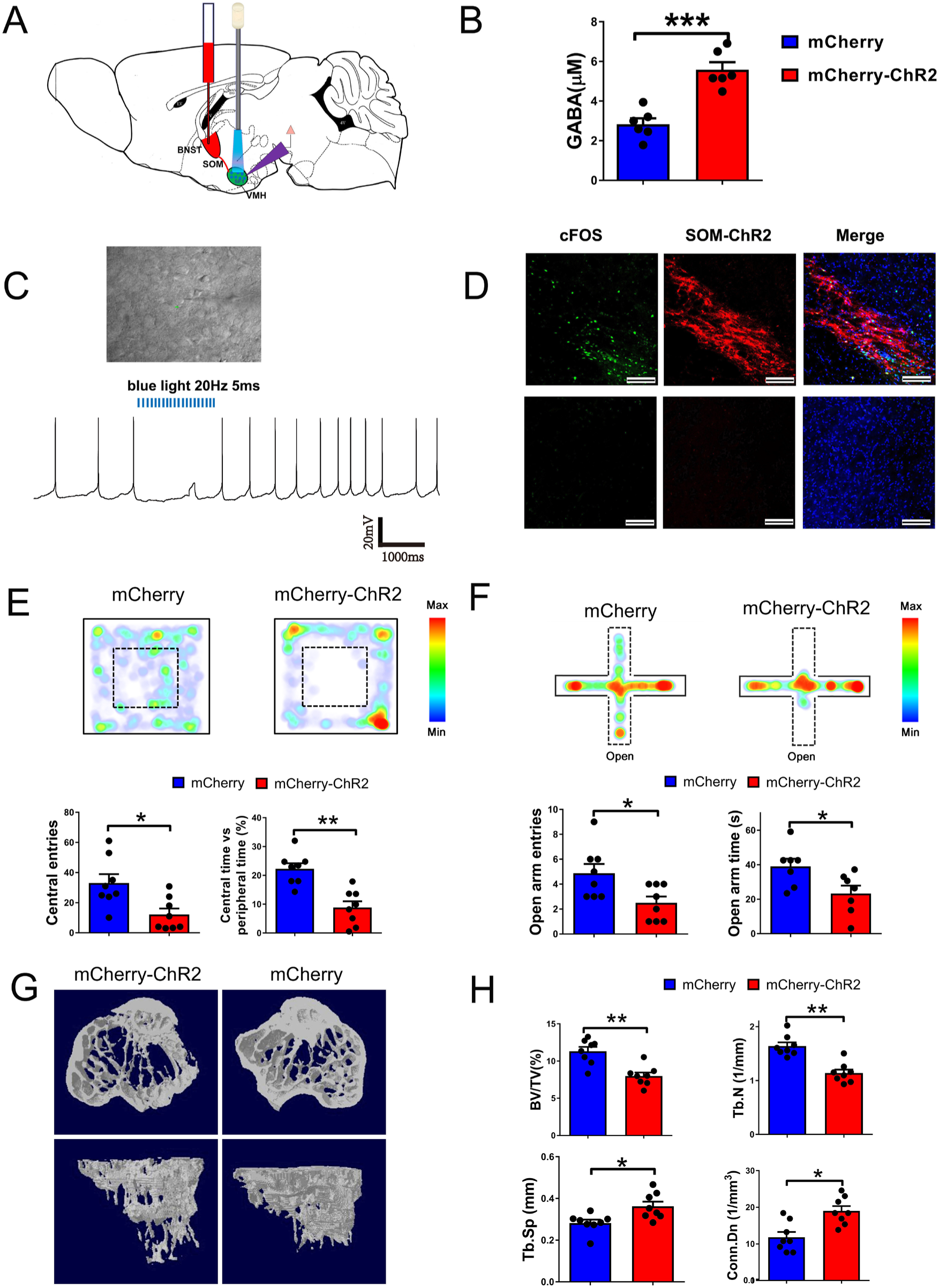
Activation of the SOM neural projection from the BNST to the VMH triggered bone loss in the absence of stressors. **(A)** Schematic showing AAV-Ef1α-DIO-ChR2-mcherry virus injected into the BSTLP region; blue light was used to illuminate the VMHdm region containing SOM neural terminals. **(B)** Quantification of the GABA level in SOM-ChR2 groups and SOM-eYFP group; values represent mean±SEM (n=6 per group; \*\*\**p*<0.001; Student’s t test). **(C)** Representative electrophysiological recordings of the inhibited firing of SF1 neurons after stimulating the VMHdm with light in SOM-ChR2 mice. **(D)** c-fos staining of the SOM positive neurons in mCherry-ChR2 group (top panel) and control (bottom panel) after blue light stimulation; scale bar, 100 μm. **(E)** Open field test comparing mCherry-ChR2 and mCherry groups; mice in mCherry-ChR2 group spent less time and made fewer entries into the central area; values represent mean±SEM (n=8 per group; \**p*<0.05; \*\**p*<0.01; Student’s t test). **(F)** EPM test comparing mCherry-ChR2 and mCherry groups; mCherry-ChR2 mice exhibited significantly less entries and spent less time into the open arms; values represent mean±SEM (n=8 per group; \**p*<0.05; Student’s t test). **(G)** Micro-CT analysis of bone structure of mCherry-ChR2 and mCherry groups 4 wks after light stimulation began; a significantly low bone mass phenotype was observed in the mCherry-ChR2 group compared to the mCherry group. **(H)** MicroCT analysis of parameters including trabecular bone volume/tissue volume (BV/TV), the trabecular number (TbN), trabecular separation (Tb.Sp), and connectivity density (Conn. Dn) in mCherry-ChR2 and mCherry groups; values represent mean±SEM (n=8 per group; \**p*<0.05; \*\**p*<0.01; Student’s t test).

### Inhibiting the activity of SOM neurons arrests anxiety-induced bone loss

To further determine the indispensability of this population of SOM neurons in anxiety-induced bone loss, we first observed enhanced c-fos signals in localized SOM neurons in the anxiety group (Figure 6B), and that some SOM neurons were stained by anti-corticotrophin releasing factor (CRF) antibody (Figure 6C). We then established an anxiety-induced bone loss model with SOM-Cre mice, and selectively silenced the SOM neurons using the DREADD (Designer Receptor Exclusively Activated by Designer Drugs) technique (Figure 6A). Briefly, AAV-Ef1α-DIO-hM4Di-mcherry and AAV-Ef1α-DIO-mcherry viruses were injected into SOM-Cre mice and Clozapine-N-oxide (CNO) was administered later during testing (Figure 6D). Electrophysiological recordings showed that somatostatin neuronal activity was selectively inhibited by CNO (Figure S3F and S3G). Anxiety levels were assessed at eight weeks and OF tests showed that mice in mCherry-hM4Di group had more central area entries and spent more time in central region compared to the mCherry group (Figure 6E) and EPM tests showed that the mCherry-hM4Di group had significantly more entries and spent more time into the open arms compared with the mCherry group (Figure 6F). Importantly, bone mass and structure were significantly better in the mCherry-hM4Di group compared to the mCherry group (Figure 6G): 25% higher BV/TV, 20% higher TbN, and 15% lower Tb.Sp (Figure 6H). The mCherry-hM4Di group had lower NE levels compared with the mCherry group, which was statistically significant (Figure S3H). Together, the DREADD experiments demonstrated that the specific population of SOM neurons in the BSTLP region were indispensable for driving anxiety-like behaviors and induced bone loss, and inhibition of the SOM neuron population was able to arrest anxiety-induced bone loss.

**Fig. 6.**
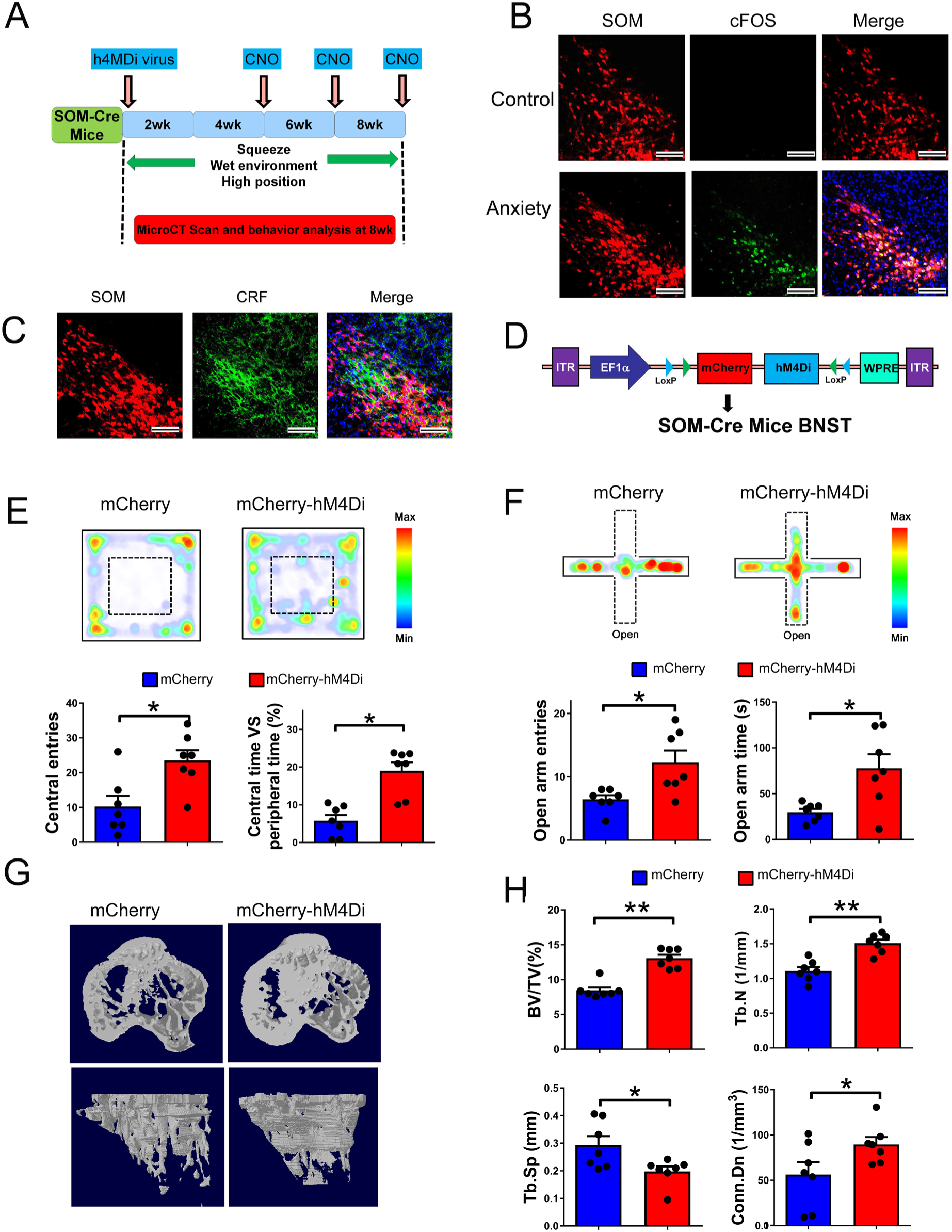
Somatostatin neurons in the BNST are indispensable for anxiety-induced bone loss. **(A)** Schematic showing the silencing SOM neurons using DREADD (Designer Receptor Exclusively Activated by Designer Drugs) technique in anxiety-induced bone loss model; clozapine-N-oxide (CNO) was used at 4, 6 and 8 wks. **(B)** c-fos staining of somatostatin positive cells in anxiety and control groups; enhanced positive signals were observed in the anxiety group; scale bar, 50 μm. **(C)** Immunostaining of corticotrophin releasing factor (CRF) on somatostatin positive neurons in anxiety-induced bone loss model with SOM-Cre mice; scale bar, 50 μm. **(D)** Schematic showing AAV-Ef1α-DIO-hM4Di-mcherry virus injection into the BNST region of SOM-Cre mice. **(E)** Open field test comparing mCherry and mCherry-hM4Di groups; mice in mCherry-hM4Di group exhibited more central entries and spent more time in central area; values represent mean ± SEM (n=7 per group; \**p*<0.05; Student’s t test). **(F)** EPM test comparing mCherry and mCherry-hM4Di groups; the mCherry-hM4Di group exhibited significantly more entries and spent more time into the open arms; values represent mean±SEM (n=7 per group; \**p*<0.05; Student’s t test). **(G)** Micro-CT analysis of bone structure comparing mCherry and mCherry-hM4Di groups after inhibition of SOM neurons; the reduced bone mass phenotype was arrested in mCherry+hM4Di group. **(H)** MicroCT analysis of the parameters including trabecular bone volume/tissue volume (BV/TV), the trabecular number (TbN), trabecular separation (Tb.Sp), and connectivity density (Conn. Dn) in mCherry group and mCherry-hM4Di group; values represent mean±SEM (n=7 per group; \**p*<0.05; \*\**p*<0.01; Student’s t test).

### NTS is downstream of the VMHdm and necessary for bone loss induced by SOM neurons

Stimulation of SOM neural projections from the BNST to the VMHdm resulted in bone loss in the absence of stressors, and SOM neurons were indispensable for anxiety-induced bone loss. We were next interested in determining the downstream effectors of this neural circuit. SF1 neurons send neural projections to regulate autonomous activities (*27*), and we injected retrograde virus into the murine trabecular bone of tibia to retrogradely label the neural circuitry innervating the bone (Figure S6A). Besides the VMH and the BNST, we observed strong signals in the nucleus tractus solitaries (NTS), which is known to modulate sympathetic activity (*28, 29*) (Figure S6B). Then we injected AAV-Ef1α-DIO-mcherry virus precisely into the VMHdm region of SF1-Cre mice (Figure 7A) and observed, interestingly, that SF1 neurons did indeed send neural projections to the medial solitary nucleus (SolM; Figure 7B), which is a subregion of NTS that mainly contains vlgut2 neurons (Figure 7C). We then confirmed the BNST-VMH-NTS neural pathway using a TRIO experiment (viral-genetic tracing of the input-output organization of the central neural circuit) (Figure 7D), and SF1 neurons that were innervated by upstream BNST neurons did indeed send direct neural projections to the NTS region (Figure 7E). To further confirm the function of localized downstream Vglut2 neurons in bone loss, we selectively inhibited those Vglut2 neurons in NTS specifically innervated by SF1 neurons using the DREADD technique (Figure 7F); the HM4Di gene was selectively expressed in SolM (Figure S6D) and inhibition of Vglut2 neurons resulted in higher NE levels in the HM4Di group compared to the eYFP group (Figure S6E). We used a micro-CT scan to analyze bone structure following chemogenetic inhibition of Vglut2 neurons and compared to the eYFP group, the hM4Di group had significantly lower bone mass and bone structure (Figure 7G): 15% lower BV/TV, 18% lower trabecular number (TbN) and 10% higher TbSp (Figure 7H). These data suggested that the vglut2 neurons in SolM were specifically employed by SF1 neurons to regulate anxiety-induced bone loss.

**Fig. 7.**
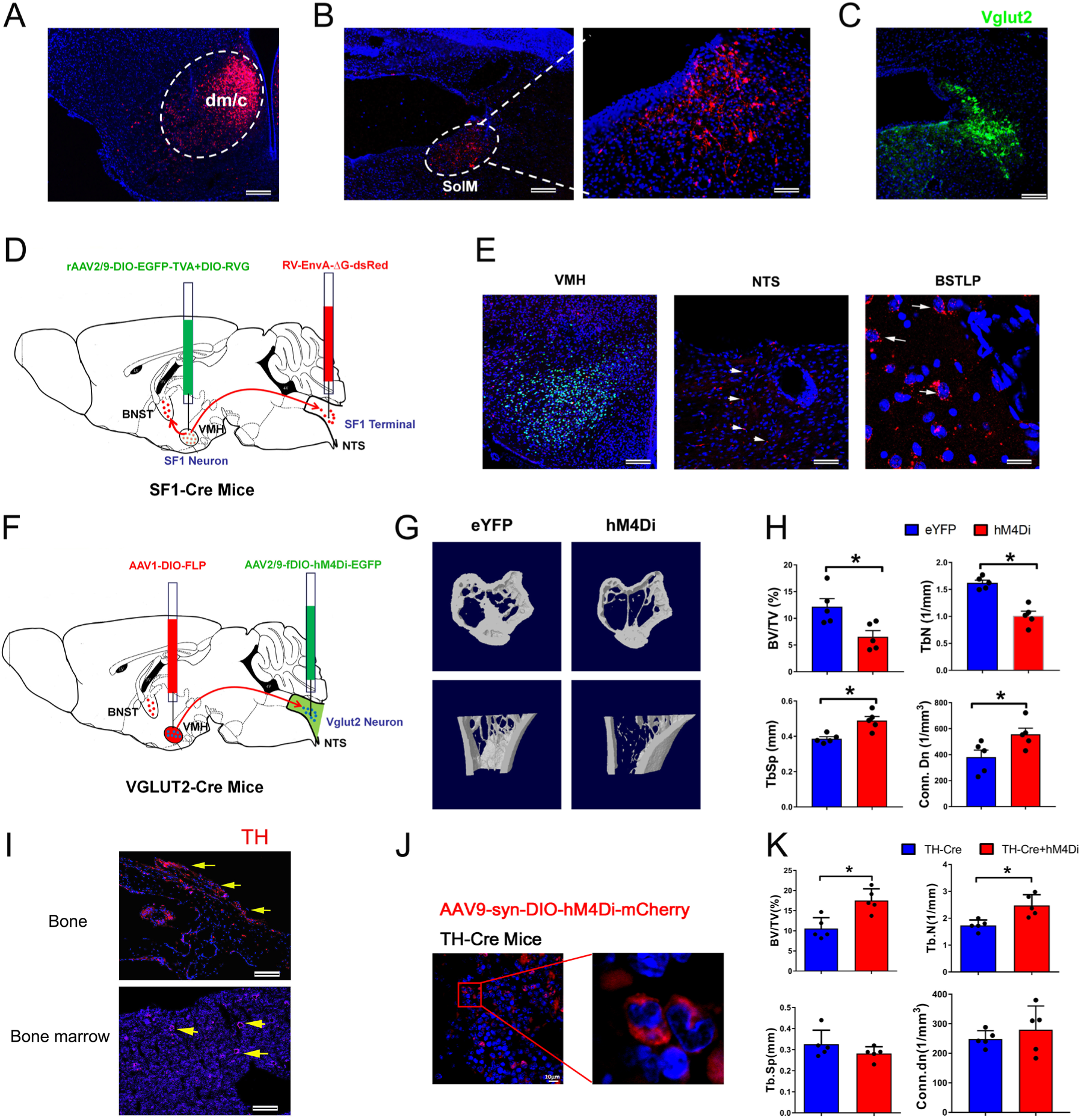
The NTS and the sympathetic system were downstream of the SOM neuron-mediated anxiety-induced bone loss. **(A)** Representative images of the VMH region with the virus expressing mCherry; scale bar, 100 μm. **(B)** Representative image of neural projections in the SolM, a subregion of the NTS, receiving projections from SF1 neurons in the VMH; scale bar, 100 μm (left); 50 μm (right). **(C)** Immunostaining of vGlut2 in SolM region of the NTS, scale bar, 50 μm. **(D)** Schematic showing the viral genetic tracing of the connection of BNST−VMH−NTS pathway using the TRIO method; AAV2-DIO-EGFP-TVA+DIO-RVG virus was injected into VMH region and RV-EnvA-ΔG-dsRed virus was injected into the NTS of the SF1-Cre mice. **(E)** Representative images showing the labeled SF1 neurons in VMH, SF1-projecting neural terminals in NTS (arrowheads) and input neurons in BNSTLP region (arrow). **(F)** Schematic showing the silencing of the Vglut2 neurons innervated by SF1 neurons using DREADD technique; AAV1-DIO-FLP virus injected into VMH region and AAV-fDIO-hM4Di-eYFP virus injected into the NTS of the Vglut2-Cre mice. **(G)** Micro-CT analysis of bone structure of eYFP group and eYFP-hM4Di group after inhibiting the Vglut2 neurons; a significantly low bone mass phenotype was observed in hM4Di group. **(H)** MicroCT analysis of the parameters including trabecular bone volume/tissue volume (BV/TV), the trabecular number (TbN), trabecular separation (Tb.Sp), and connectivity density (Conn. Dn) in eYFP and hM4Di groups; values represent mean±SEM (n=5 per group; \**p*<0.05; Student’s t test). **(I)** Immunostaining of tyrosine hydroxylase (TH) in bone and bone marrow; positive signals were observed in both the bone and the bone marrow (arrow); scale bar, 100 μm. **(J)** The expression of hM4Di-mCherry on TH-positive nerve fibers in bone marrow of TH-Cre mice. **(K)** MicroCT analysis of the parameters including trabecular bone volume/tissue volume (BV/TV), the trabecular number (TbN), trabecular separation (Tb.Sp), and connectivity density (Conn. Dn) in TH-Cre and TH-Cre+hM4Di groups; values represent mean± SEM (n=5 per group; *p<0.05; Student’s t test).

Immunostaining showed that tyrosine hydroxylase (TH), a marker for sympathetic nerve axons, could be identified both in tibia bone and bone marrow (Figure 7I). To further investigate the effects of NE on the osteogenic differentiation process, we cultured osteoprogenitor cells from murine bone marrow and found that beta-2 and beta-3 adrenergic receptors were positively expressed in osteoprogenitor cells (Figure S6F). Quantification of ALP showed that osteoblastic differentiation of osteoprogenitors was significantly inhibited by NE compared to the control group (Figure S6G). RT-PCR analysis showed that the normalized expression of *Runx2*, *Alp*, *Col1a1*, and *Opn* in the NE group were significantly lower than those of the control group at two weeks of osteogenic differentiation (Figure S6H); we also confirmed the inhibitory effects of NE on osteogenesis *in vivo* by performing gene expression analysis with freshly isolated bone marrow cells and found that the expression of *Runx2*, *Alp*, *Col1a1*, and *Opn* was lower in the anxiety group compared with the control group; however, treatment using beta-blockers significantly elevated the expression of these osteogenesis-related genes (Figure S6I). Finally, we used the DREADD technique to selectively silence the TH-positive nerve fibers in bone marrow of TH-Cre mice (Figure 7J). The hM4Di group had significantly higher bone mass compared to the TH-Cre control group: 35% higher BV/TV, 29% higher trabecular number (TbN) (Figure 7K). These data consistently demonstrate that tyrosine hydroxylase in the sympathetic system and NE relay the BNST-VMH-NTS neural pathway to modulate the anxiety induced bone loss.

## Discussion

The brain plays an essential role in the integration of cognitive and emotional neural information to regulate food intake and to maintain peripheral metabolic homeostasis. Recent studies have discovered important evidence supporting the close association between bone metabolism disturbance and psychiatric disorders, such as anxiety (*1–3*), however, the neural mechanism behind this was previously unclear. Our study is the first to provide experimental evidence that the neural circuitry BNST-VMH-NTS can regulate anxiety-induced bone loss. We identified a specific group of somatostatin neurons in the posterior region of BNST projecting to the VMH that are responsible for anxiety-induced bone loss. These results not only indicate that the GABAergic neural circuitry from the forebrain to the hypothalamus can exert a fundamental influence on the homeostasis of bone metabolism, but they also provide a new central target for therapeutic intervention of anxiety-induced bone metabolism disorders.

### Anxiety-related disorders, central molecules and neural circuitry regulating bone metabolism

Recent studies have suggested that mood disorders are closely related with bone metabolism abnormalities. For example, bone quality is lower amongst men and women with a history of mood disorders (*5*), older men with probable anxiety have a greater than four-fold risk of experiencing a fracture (*4*) and bone development is compromised in children with the autism spectrum disorders (*30*). Psychological changes due to extreme environments, such as prolonged, simulated spaceflight can also lead to bone metabolism disorders (*9–11*). Whilst microgravity is an established cause of bone loss during the spaceflight, decreased bone formation in astronauts progressively deteriorates under normal gravity after returning to earth (*9*). In this study we collaborated with the China Astronaut Research and Training Center to perform a longitudinal study of chronic stress on anxiety and bone loss of crewmembers under normal gravity. We observed significantly raised anxiety scores and continuous elevated levels of norepinephrine (NE) during the period of chronic stress. Decreased BMD and a continuous reduced level of bone formation markers also reflected reduced bone formation during the experiment. Although there were only four participants, our longitudinal data demonstrated consistently that chronic stress in crewmembers indeed resulted in both decreased bone density and elevated anxiety levels under normal gravity.

Central molecules and neuropeptides can play important roles in the regulation of bone metabolism (*31*). Leptin inhibits bone formation through the sympathetic nervous system (*14*) and orexin is a critical regulator of skeletal homeostasis and exerts dual regulation of bone mass (*16*). Specific neuronal subtypes and circuits can also exert influence on the regulation of bone mass (*32*), such as AGRP neurons in the arcuate nucleus that regulate bone mass through the sympathetic nervous system (*33*). While genetic knocking down of CamKKβ and CamKIV in SF1 neurons in the VMH leads to severe low-bone mass (*21*), serotonin neurons in the raphe nucleus send axonal projections to VMH SF1 neurons and promote bone mass through Htr2c receptors on the SF1 neurons (*15*). These genetic and molecular studies provide useful information of neural molecules regulating bone mass; however, up until now, studies attempting to dissect the central neural regulation of bone at the circuit level have been scarce and the specific neural circuits regulating anxiety and bone mass not clear. We used an established mouse model to dissect the neural circuitry regulating this anxiety-induced bone loss and identified a neural circuit from the forebrain to the hypothalamus that relayed anxiety information and regulate bone mass. Specifically, GABAergic neural projections in the VMH can regulate anxiety-like behavior and reduce bone mass. Our systematic study using both human participants and an animal model thus aids further understanding of the neural mechanism of anxiety-induced bone loss.

### The VMH, an important hypothalamic region, regulates both behavior and bone metabolism

The VMH is heavily involved in regulating energy metabolism and glucose homeostasis (*34–36*). For example, leptin can directly activate SF1 neurons in the VMH to regulate normal body weight homeostasis (*13*), the neural circuit from the PBN to the VMH controls counterregulatory responses to hypoglycemia (*20, 37*), and serotoninergic projections from the raphe nucleus to the VMH regulate SF1 neurons to promote bone accrual (*15*). In addition, AP1 alterations in SF1-expressing neurons in the VMH increase energy, but decrease bone density (*24*), suggesting that the heterogeneous or distinct neuronal circuits in VMH mediate the disparate effects of regulating bone homeostasis and energy metabolism.

The VMH also plays an important role in emotional state and the dorsomedial part of the VMH (VMHdm) is selectively involved in predator fear (*18*). Ablation of SF1 neurons in the VMHdm attenuate both innate and learned defensive behaviors (*38*), and activation of SF1 neurons in the VMHdm can induce defensive-like motor and autonomic responses (*23*). Serotonin receptors on VMH neurons not only contribute to the regulation of bone metabolism (*15*), but also in generalized anxiety (*39*). In addition to these findings, we further show that the link between stress-induced anxiety and bone regulation occur at the level of the VMHdm. Indeed, our findings show that the activation of the GABAergic neural projections in the VMHdm not only increased anxiety levels, but also reduced bone mass, thus revealing the GABAergic neural projections in the VMHdm as an important link between anxiety-like behavior and bone metabolism. This study is the first to show that convergent regulation of bone mass and anxiety is integrated in the GABAergic circuit in the VMHdm and provides evidence that psychiatric disorders, such as anxiety, can induce bone loss through GABAergic neural projections in the VMH.

### A GABAergic neural circuit involving the BNST utilizes somatostatin neurons to regulate anxiety and bone loss

The BNST is an important brain region in the regulation of stress and anxiety (*26*). Specific excitatory and inhibitory neuron types in the BNST can utilize distinct neural pathways to integrate and process aversive stimulation (*40*). The BNST also serves as a key relay connecting forebrain structures to hypothalamic and brainstem regions that regulate feeding, autonomic and neuroendocrine functions (*41, 42*). The GABAergic inhibitory synaptic input from BNST to the lateral hypothalamus (LH) can suppress local glutamatergic neurons to control food intake (*43*). In the hypothalamus, both local and long range GABAergic projections play important roles in regulating behaviors and metabolism (*44–46*). Our electrophysiological recordings and viral tracing methods show that the activation of GABAergic neural projections from the BNST to the VMHdm can release GABA to inhibit SF1 neurons; this long range GABAergic axonal projection plays an important role in regulating both anxiety-like behavior and bone loss.

A specific population of SOM neurons in the lateral posterior region of the BNST was identified that sends GABAergic neural projections to the VMH. Importantly, the specific activation of the GABAergic output of these SOM neurons induced both anxiety-like behavior and bone loss in the absence of stressors. Somatostatin-expressing GABAergic neurons constitute a major class of inhibitory neurons in different regions of the brain (*47*) and are important in anxiety and neuroendocrine function. Enhanced activity of SOM neurons in the dorsal BNST drives elevation of anxiety (*48*), global dysfunction of SOM neurons contributes to altered neuroendocrine function and to aberrant inhibitory neurotransmission (*49*), and SOM neurons in the hypothalamic arcuate nucleus and tuberal nucleus control energy balance and feeding (*50, 51*). In this study we found that activation of a newly discovered population of SOM neurons in lateral posterior BNST led to decreased bone mass, and that inhibition of this population of neurons coupled with anxiety led to lower anxiety and bone loss than that observed without the neural inhibition. These data systematically reveal how SOM neurons in a specific BNST region and its GABAergic output is both indispensable for and able to drive anxiety-induced bone loss.

### The NTS and sympathetic nervous system as a central relay to regulate bone mass

Periosteum, cortical bone and bone marrow are all richly innervated by both sympathetic and sensory nerve fibers (*52, 53*). Using virus-based transneuronal tracing from murine bone marrow, we found that neurons in certain brain nuclei, such as the BNST, VMH and downstream NTS was retrogradely labeled, thus providing anatomical evidence of central neural control of bone mass. As the relay of the central nervous system, both the sensory and sympathetic nervous systems are crucial in regulating bone homeostasis and metabolism (*54–56*); for example, calcitonin gene-related peptide (CGRP) expressing sensory nerves in peripheral cortical femur enhances osteogenic differentiation of osteoprogenitor cells (*57*). The sympathetic nervous system plays important roles in the regulation of bone metabolism (*14*), and sympathetic tone signals in osteoblasts can inhibit CREB phosphorylation to decrease osteoblast proliferation (*54*). Our study shows that the sympathetic neurotransmitter, norepinephrine (NE) was significantly higher in the anxiety and SOM-ChR2 groups, suggesting elevated sympathetic tone following activation of the BNST-VMH neural projections.

We observed that SF1 neurons send direct axonal projections to a specific region of NTS, which regulates the activity of the sympathetic nervous system. To further investigate the role of NTS in relaying the information, we used a TRIO experiment to confirm the BNST-VMH-NTS neural pathway; we selectively inhibited Vglut2 neurons in NTS that are specifically innervated by SF1 neurons using the DREADD technique to confirm the function of Vglut2 neurons in the NTS during anxiety-induced bone loss. Higher levels of NE and lower bone mass and bone structure in the HM4Di group suggests that vglut2 neurons in the NTS were indeed employed by SF1 neurons to mediate anxiety-induced bone loss. Our data thus verified that the NTS is downstream of the BNST-VMH circuitry and necessary for SOM neuron-induced bone loss. Along with the increased bone density after specifically inhibiting TH-positive nerve fibers in bone, our data consistently suggests that the activation of BNST-VMH neural projections can induce bone loss through the sympathetic nervous system and NE.

In summary, we have uncovered a new GABAergic neural circuit from the forebrain to hypothalamus used in the regulation of anxiety-induced bone loss and have also revealed a population of somatostatin neurons in the posterior region of BNST that has not been implicated in bone mass regulation. Vglut2 neurons in NTS and the peripheral sympathetic system were employed by this BNST-VMH neural circuit to regulate anxiety-induced bone loss. Our findings, therefore, not only identify the underlying central neural mechanism of the anxiety induced bone loss at circuit level for the first time, but also provide a new central target for therapeutic interventions of anxiety-induced bone metabolism disorders.

## Materials and Methods

### Animals

All procedures were carried out in accordance with protocols approved by the ethics committee of the Shenzhen Institutes of Advanced Technology, Chinese Academy of Sciences. All animals were housed at 22-25 °C on a circadian cycle of 12 hour light-dark (lights on at 07:00 am and off at 07:00 pm) with ad-libitum access to food and water. All animals used in this study were male (2-6 months old). SF1-Cre, GAD2-ires-Cre, PV-Cre, SOM-Cre, CCK-Cre, Vglut2-Cre, TH-Cre and VGAT-ChR2-eYFP mice (Jackson stock no: 012462, 010802, 008069, 013044, 011086, 016963, 008601 and 014548 respectively) were obtained from Jackson Laboratories. Adult male C57BL/6 were purchased from Guangdong Medical Laboratory Animal Center, Guangzhou, China. All transgenes were used as homozygotes. Mice experiments were performed during the light cycle. No animals were involved in any previous studies and they were sacrificed during the light cycle.

### Human participants

Three male and one female participant (Chinese, aged 23-45 years) were selected as the 180-day Controlled Ecological Life Support System (CELSS) crew following consideration of psycho-physiological health, occupational background and other similar criteria. This research was conducted in accordance with the principles expressed in the Declaration of Helsinki. The ethical committee of China Astronaut Research and Training Center reviewed and approved 180-day study, and all crewmembers gave their written informed consent.

### 180-day Controlled Ecological Life Support System (CELSS) integrated experiment

A 180-day CELSS integrated experiment was implemented to simulate a space residence to prepare for future deep-space exploration. It lasted for 180 days, from Jun 17^th^ 2016 to Dec 14^th^ 2016. The experimental facility was located in the Space Institute of Southern China, Shenzhen, China. The 180-day CELSS crew lived in this spaceship-like habitat with continuous temporal and spatial isolation, realistic mission activities and other major special conditions of a space station.

Bone mineral density (BMD) was determined once prior to (Day 0) and once after (Day 180) the experiment by dual-energy X-ray absorptiometry (DXA) as previously described. The Self-Rating Anxiety Scale (SAS) was used to measure crewmember anxiety levels; this focuses on the most common general anxiety disorders. Immediately prior to and during the mission, crewmembers filled out several computerized SAS questionnaires and rating scales. There were 20 questions with 15 increasing anxiety level questions and five decreasing anxiety questions. The four crewmembers indicated their current anxiety status by responding to all questions using a 4-point scale from ‘none of the time” to “most of the time”.

Concentrations of Procollagen I carboxy-terminal propeptide (PICP) and alkaline phosphatase (ALP) in the serum were determined using the ELISA Kits (Lianshuo Biological, Shanghai, China). Cortisol, epinephrine, norepinephrine in the serum were determined using the ELISA Kits (Roche Diagnostics, USA). The experiments were carried out by Kingmed diagnostics (Guangzhou, China) according to manufacturer instructions.

### Stress induced anxiety procedures

C57BL/6 or SOM-Cre mice were exposed to environmental stressors for eight weeks. The following stressors were applied: (i) tight squeeze for two hours where four mice were housed in a relatively small box (3 cm×5 cm×7 cm). (ii) wet environment for six hours where water was added to home cages to moisten bedding to make damp without generating large pools and (iii) high position stress where mice were placed on a platform raised 100 cm above floor height for two hours. Stressors were randomized and counterbalanced such that each mouse received the same number of each stressor across consecutive days or the duration of eight weeks. Efficacy of the induced-anxiety procedures was confirmed by animal behavior studies detailed below.

### Immunostaining

Brains were fixed in 4% paraformaldehyde (PFA) at 4 °C overnight and cryosectioned at a thickness of 20 μm. Sections were then rehydrated and blocked by goat serum. The sections were incubated with primary antibodies to SF1 (1:500, Abcam, ab65815), GAD65 (1:500, Abcam, ab11070), Vglut2 (1:400, Abcam, ab79157), TH (1:500, Sigma, T8700), Parvalbumin (1:500, Millipore, MAB1572), Somatostatin (1:500, Millipore, AB5494), Cholecystokinin (1:500, Abcam, ab27441) or cfos (1:500, Cell Signaling Technology, mAb2250). The sections were then washed and labeled with fluorescence-conjugated corresponding secondary antibodies (Jackson ImmunoResearch, 111-545-003, 315-545-003). The sections were counterstained with Hoechst 33342 and then mounted for image acquisition. Images were taken under a microscope (ECLIPSE 50i, Nikon, Melville, NY, USA). The number of cfos-positive cells in the BNST and other regions were counted in eight sections (three adjacent levels) from each mouse, and each group contained three mice. The mean cell number per section was compared with the different experimental groups during data analyses.

### Microdialysis and determination of GABA levels

Mice were anesthetized with sodium pentobarbital (100 mg/kg) and placed in a stereotaxic frame. A microdialysis probe (MER-10 mm guide, 2 mm membrane, Bioanalytical Systems Inc.) was implanted into the Ventromedial Hypothalamus (VMH) region (AP=-1.58 mm; ML=±0.3 mm; DV=-5.35 mm). The probes were perfused with artificial cerebrospinal fluid (ACSF) (124 mM NaCl, 3 mM KCl, 2.4 mM CaCl2, 1.3 mM MgSO4, 10 mM glucose, and 10 mM HEPES; pH=7.3) for 90 min before collecting samples. Samples were collected for 30 min during the experiment. GABA concentrations in the dialysate were determined using an ELISA Kit (MyBioSource, Canada) and the experiments were carried out according to the manufacturers’ instructions. In brief, the optical density was determined at 450 nm using the microplate reader (Synergy 4, Biotek). The concentrations of GABA were determined, normalized to that of the control group, and compared to those from the different experimental groups.

### MicroCT analysis

The proximal tibia of mice in different experimental groups were scanned and analyzed using a SkyScan 1076 Micro-CT system and software (SkyScan, Kontich, Belgium) with a voxel size of 9 um, voltage of 50 kV, exposure time of 1018 ms, frame averaging on and beam filtration filter 1.0 mm aluminum. After scanning, knee joints were three-dimensionally reconstructed by SkyScan recon software. A cuboid of trabecular bone beneath the growth plate with size of 1.15 x 1.15 x 0.58 mm^3^ was selected. Percentage bone volume (BV/TV, %), trabecular number (TbN, 1/mm), trabecular separation (TbSp, mm), and connectivity density (1/mm^3^) were calculated for the tibia trabecular bone using CT-Analysis with thresholding of 60-255. Additionally, diagrammatic images were generated by CT-Volume with a well-established protocol (*58*).

### Histological staining

Proximal tibia from different groups were isolated at eight weeks after light stimulation. Tissue processing and sectioning was carried out as previously described (*59*). In brief, tissue samples were fixed in 4% PFA for 48 hours at 4 °C and then decalcified in 4% ethylenediaminetetraacetic acid (EDTA) for 30 days. Following decalcification, tibia were embedded in paraffin and sectioned at 5 μm thickness. Hematoxylin & Eosin (HE) staining were performed separately on consecutive tissue sections and images were taken using a microscope (ECLIPSE 50i, Nikon, Japan). For immunostaining, cryosections of tibia were incubated with primary TH antibody (1:500, Sigma, T8700) for 60 min and then washed and labeled with fluorescent secondary antibody (Jackson ImmunoResearch, 111-545-003, 315-545-003). Sections were counterstained with Hoechst 33342 and then mounted for image acquisition.

### Trans-synaptic tracer labeling

All animal procedures were performed in Biosafety level 2 (BSL2) animal facilities. To determine whether the BNST-VMH pathway was innervated by GABAergic neurons in the BNST, SF1-Cre mice (20–25 g) were used for transmono-synaptic tracing based on the modified rabies virus. First, a mixture of AAV2/9-EF1a-FLEX-TVA-GFP and AAV2/9-EF1a-Dio-RV-G (1:1, total volume of 200 nl) was stereotaxically injected into the VMH region using the following coordinates (AP=-1.58 mm; ML=±0.3 mm; DV=-5.35 mm). Mice were housed on a regular 12-hour light/dark cycle with food and water ad-libitum during recovery. Three weeks later, 200 nl of EnvA-pseudotyped rabies virus (EnvA-RV-DsRed) was injected into the VMH using the previously defined coordinates.

To verify the connectivity of BNST-VMH-NTS pathway, we modified the mono-synaptic rabies tracing strategy (*60*). On the first day, we injected a mixture of 200 nl AAV2/9-EF1a-Dio-histone-TVA-GFP and AAV2/9-EF1a-Dio-RV-G (1:1) into the VMH of SF-1-Cre mice. Three weeks later, to allow the accumulation of SF1 neuronal TVA to be transported to axon terminals in the NTS (AP=-7.32 mm; ML=0.3 mm; and DV=-4.38 mm), we injected 200 nl of EnvA-RV-dsRed into the NTS of these mice. Thus, we specifically infected NTS-projecting SF1 neurons and traced their inputs in BNST. Mice were sacrificed 2 weeks after RV injection.

### Virus injection and light stimulation

Mice were anesthetized with pentobarbital sodium (i.p., 100 mg/kg), and then fixed in a stereotaxic apparatus (RWD, China). During surgery and virus injections, mice were kept anesthetized with isoflurane (1%). The skull above targeted areas was thinned with dental drill. A microsyringe pump (UMP3/Micro4, USA) was employed to conduct the virus injections with a 10 μl syringe connected to a 33-G needle (Neuros; Hamilton, Reno, USA). For optogenetic activation, 300 nl AAV5-DIO-ChR2-mCherry (10^9^ TU/ml) or AAV5-DIO-mCherry (10^9^ TU/ml) was injected into the BNST region (AP, −0.22 mm, ML, ±0.75 mm, DV, −4.5 mm) of SOM-Cre mice; mice were housed for four to six weeks following injection for viral expression before initiation of experiments.

For optical terminal stimulation, custom-made optic fiber cannulae (200 um diameter, 0.37 NA. fiber with 1.25 mm ceramic ferrule; NEWDOON, Hangzhou) were implanted unilaterally above the VMH (AP, –1.58 mm; ML, ±0.3 mm; DV, −5.35 mm) of SOM-Cre mice four weeks after the BNST virus injection. Dental cement was applied to cover the exposed skull completely and to secure the implant. Then blue light stimulation (470 nm) was performed at 20 Hz with intervals of 5 min for a total of 30 min in the mCherry and mCherry-ChR2 groups. The light stimulation was conducted every three days and lasted for eight weeks in mCherry and mCherry-ChR2 groups. The anxiety behavior tests and bone analysis were performed at the end of the paradigm. After the final light stimulation mice were perfused with 4% PFA and brain tissues were removed for immunostaining analysis.

### Chemogenetic inhibition of neurons

For chemogenetic inhibition of the somatostatin neurons in BNST, AAV5-DIO-hM4Di-mCherry (10^9^ TU/ml) or AAV-DIO-mCherry (10^9^ TU/ml) was injected bilaterally into the BNST (AP=-0.22 mm, ML=±0.75 mm, DV=4.5 mm) of SOM-Cre mice. Mice were housed for four weeks following injection for viral expression before initiation of experiments. Clozapine N-oxide (CNO) (1 mg/kg, sigma, C0832) or saline was delivered by intraperitoneal injection. Control saline injections contained an equivalent amount of Dimethyl sulfoxide (DMSO) (0.6%).

To selectively manipulate Vglut-2 neurons in NTS innervated by SF1 neurons, we first bilaterally injected AAV1-Ef1α-DIO-FLP (300 nl) into VMH (AP=-1.58 mm; ML=±0.3 mm; and DV=-5.35 mm). Then, AAV-fDIO-hM4Di-eYFP virus was injected into the NTS (AP=-7.32 mm; ML=0.3 mm; and DV=-4.38 mm) during the same surgery. After completion of the injection, the needle was kept for 10 min for virus diffusion purposes and then slowly withdrawn. Following surgery, mice were allowed to recover from anesthesia under a heat lamp. Lincomycin hydrochloride and lidocaine hydrochloride gel was applied to the sterilized incision site as an analgesic and anti-inflammatory drug. Mice were allowed to recover for two to three weeks after surgery.

To investigate the role of TH-positive neural terminals in bone metabolism, we injected 500 nl AAV2/9-DIO-hM4Di-mCherry or AAV2/9-DIO-mCherry in bone marrow cavity of TH-Cre mice to selectively label local TH terminals. Six weeks after the virus injection, CNO (1mg/kg) was delivered by intraperitoneal injection every three days for four weeks. At the end of the stimulation, Micro-CT scanning and analysis were performed on the collected samples.

### Electrophysiology

Procedures for preparing acute brain slices and performing whole-cell recordings with optogenetic stimulations were similar to those described previously. Coronal slices (300–400 μM) were prepared using a vibratome (VT-1000S, Leica) in an ice-cold choline-based solution containing 110 mM choline chloride, 2.5 mM KCl, 0.5 mM CaCl2, 7 mM MgCl2, 1.3 mM NaH2PO4, 1.3 mM Na-ascorbate, 0.6 mM Na-pyruvate, 20 mM glucose and 2.5 NaHCO3, saturated with 95% O_2_ and 5% CO_2_. Slices were incubated in 32 °C oxygenated artificial cerebrospinal fluid (125 mM NaCl, 2.5 mM KCl, 2 mM CaCl2, 1.3 mM MgCl2, 1.3 mM NaH2PO4, 1.3 mM Na-ascorbate, 0.6 mM Na-pyruvate, 10 mM glucose and 2.5 mM NaHCO3) for at least 1 h before recording. Slices were transferred to a recording chamber and superfused with 2 ml/min artificial cerebrospinal fluid. Patch pipettes (4–7 MΩ) pulled from borosilicate glass (PG10150-4, World Precision Instruments) were filled with internal solution containing 35 mM K-gluconate, 10 mM HEPES, 0.2 mM EGTA, 5 mM QX-314, 2 mM Mg-ATP, 0.1 mM Na-GTP, 8 mM Nacl, at 280∼290 mOsmkg^−1^ and adjusted to pH 7.3 with KOH. Whole-cell voltage-clamp recording of SF1 neurons was performed at room temperature (22–25 °C) with a Multiclamp 700B amplifier and a Digidata 1440A (Molecular Devices). Data were sampled at 10 kHz and analyzed with pClamp10 (Molecular Devices) or MATLAB (MathWorks).

The eYFP-expressing SF-1 neurons in the VMH were visualized using an upright fluorescent microscope (Nikon FN-S2N). Blue light from a DG4 (Lambda, Sutter Instrument Company) (470 nm) controlled by digital commands from a Digidata 1440A was coupled to the microscope with an adaptor to deliver photo stimulation. To record light-evoked IPSCs, 20 s of 0.5–2 mW blue light was delivered through the objective to illuminate the entire field of view. The experiment was performed in the presence of AMPA receptor antagonist NBQX (50 μM) and NMDA receptor antagonist AP5 (50 μM). Both the frequency and amplitude of IPSCs were analyzed in the presence of bicuculine (50 μM).

### Gene expression analysis using RT-PCR

RT-PCR was conducted using a Light Cycler VR 480 (Roche, Switzerland) according to standard Taqman or SYBR® Green technology procedure employing fluorescence monitoring. For single cell RT-PCR, at the end of each recording, cytoplasm was aspirated into the patch pipette, expelled into a PCR tube. The single cell RT-PCR protocol was designed to detect the presence of mRNAs coding for SF-1, vGlut2, VGAT and 18S. Commercialized predesigned gene-specific primers and probes (Applied Biosystems) for *SF-1* (Mm00446826_m1), *vGlut2* (Mm00499876_m1), *VGAT* (Mm00494138_m1), and 18S rRNA control reagents (Mm03928990_g1) were obtained from ThermoFisher Scientific, and we used Single Cell-to-CTTM Kit (ambion, ThermoFisher Scientific) at a final reaction volume of 50 μl/well in 96-well plates. All procedures were conducted according to the manufacturer’s protocol.

For bone marrow mesenchymal stem cells differentiation analysis, reverse transcription and PCR amplification were performed using the ReverTra Ace qPCR RT kit (TOYOBO, FSQ-101) and SYBR® Green Realtime PCR Master Mix (TOYOBO, QPK-201), respectively. Primers used for *Runx2* were: 5’GGGCACAAGTTCTATCTGGAAAA3’, 5’CGTGTCACTGCGCTGAA3’ (Final product 71 bp); *ALP*: 5’GCCCTCTCCAAGACATATA3’, 5’CCATGATCACGTCGATATCC3’ (Final product 373 bp); *OPN*: 5’GAAACTCTTCCAAGCAATTC3’, 5’GGACTAGCTTGTCCTTGTGG3’ (Final product 589 bp); *Col1a1*: 5’GGTGAACAGGGTGTTCCTGG3’, 5’TTCGCACCAGGTTGGCCATC3’ (Final product 503 bp). *Gapdh:* 5’GCATGGCCTTCCGTGTTC3’, 5’CCTGCTTCACCACCTTCTTGAT3’ (Final product 105 bp) was used as an internal control to normalize RNA content. The experiments were carried out according to the manufacturers’ instructions.

### Animal behavior studies

The elevated plus maze test and open-field test are widely used behavioral assays for measuring anxiety in rodents. The elevated plus maze was made of plastic and consisted of two white open arms (25 × 5 cm) and two white enclosed arms (25 × 5 × 15 cm) extending from a central platform (5 × 4 cm) at 90° to form a plus shape. The maze was placed 65 cm above the floor. Mice were individually placed in the center, with their heads facing a closed arm. The number of entries and amount of time spent in the same type of arms were observed during 8 min sessions. The open-field test consisted of an 8 min session in the open-field chamber (50 ×50 cm), which was made of plastic and was conceptually divided into a central field (center, 25 × 25 cm) and a peripheral field for analysis purposes. Each individual mouse from the different experimental groups was placed in the peripheral field at the start of the test. Behaviors were recorded on video during the trial and the ANY-maze video tracking system (Stoelting, USA) was used for analysis.

### Statistical analysis

Experiment data are expressed as the mean SEM of the number of tests stated. Statistical comparisons were made using either Student’s t-test or analysis of variance (ANOVA) followed by Bonferroni multiple comparisons post-hoc test, as indicated in the figure legends. All of the statistical tests were performed using the Statview (version 10.0, SPSS, Chicago, IL) program package or Prism 7.0 software. A p-value of <0.05 was considered statistically significant.

## Supplementary Materials

Fig. S1. Biochemical parameters of mice in stress and control groups

Fig. S2. Activation of GABAergic projections in the VMHdm inhibit the firing of SF1 neurons

Fig. S3. Somatostain neurons in the BNST send GABAergic projections to the VMHdm region

Fig. S4. Activation of parvalbumin neural projections in the VMH did not induce anxiety-like behavior or bone loss

Fig. S5. Activation of the Cholecystokinin neural projection in the VMH did not induce anxiety-like behavior or bone loss

Fig. S6. Sympathetic system and NE-mediated BNST-VMH-NTS neural circuitry-induced bone loss

Fig. S7. Schematic showing that the neural circuit BNST-VMH-NTS can regulate anxiety-induced bone loss

## Acknowledgments

We are grateful to all the members of the Wang and Yang laboratories for providing comments and advice throughout the project.

## Author contributions

F.Y; Y.L; L.W designed the experiments, analyzed data and wrote the manuscript. Y.L; S.C and Z.D performed the experiments with help of G.D; D.Y; Y.W; T.W; L.Z and W.W. S.C performed electrophysiological recordings in brain slices. Y.L performed surgery and optogenetic experiments. F.Y; Y.L and L.W conceived and supervised the project.

## Data and materials availability

All data associated with this study are present in the paper or the Supplementary Materials

## Supplementary Materials

**Fig. S1.**
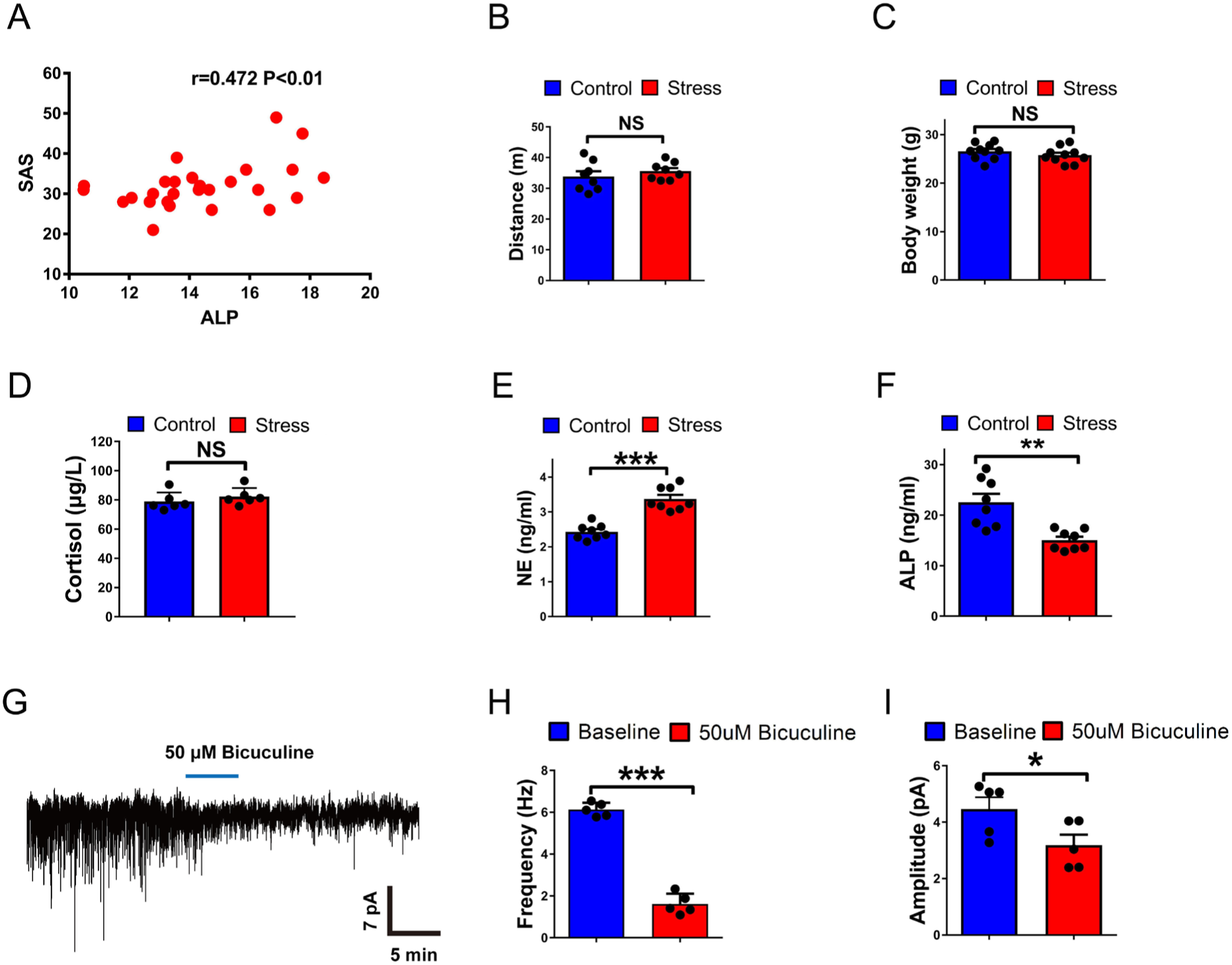
Biochemical parameters of mice in stress and control groups. **(A)** Correlation analysis using alkaline phosphatase (ALP) and Self-Rating Anxiety Scale (SAS) scores that reflect the level of anxiety. **(B)** Total distance traveled in open field test for stress and control groups; values represent mean±SEM (n=8 per group; NS, not significant; Student’s t test). **(C)** Bodyweight of mice in the stress and control groups; values represent mean±SEM (n=8 per group; NS, not significant; Student’s t test). **(D)** Cortisol levels in stress and control groups; values represent mean ± SEM (n=6 per group; NS, not significant; Student’s t test). **(E)** Quantification of the norepinephrine (NE) of stress and control groups; values represent mean± SEM (n=8 per group; \*\*\**p*<0.001; Student’s t test). **(F)** Quantification of alkaline phosphatase (ALP) of stress and control groups; values represent mean±SEM (n=8 per group; \*\**p*<0.05; Student’s t test). **(G)** IPSCs in the stress group was completely blocked by bicuculline (50 μM). **(H)** Bicuculline (50 μM) blocked the increased frequency and amplitude of IPSCs in the stress group; values represent mean ± SEM (n=5 per group; \*\*\**p*<0.001; Student’s t test). **(I)** Bicuculline (50 μM) blocked the increased frequency and amplitude of IPSCs in the stress group; values represent mean±SEM (n=5 per group; \*\*\**p*<0.001; Student’s t test).

**Fig. S2.**
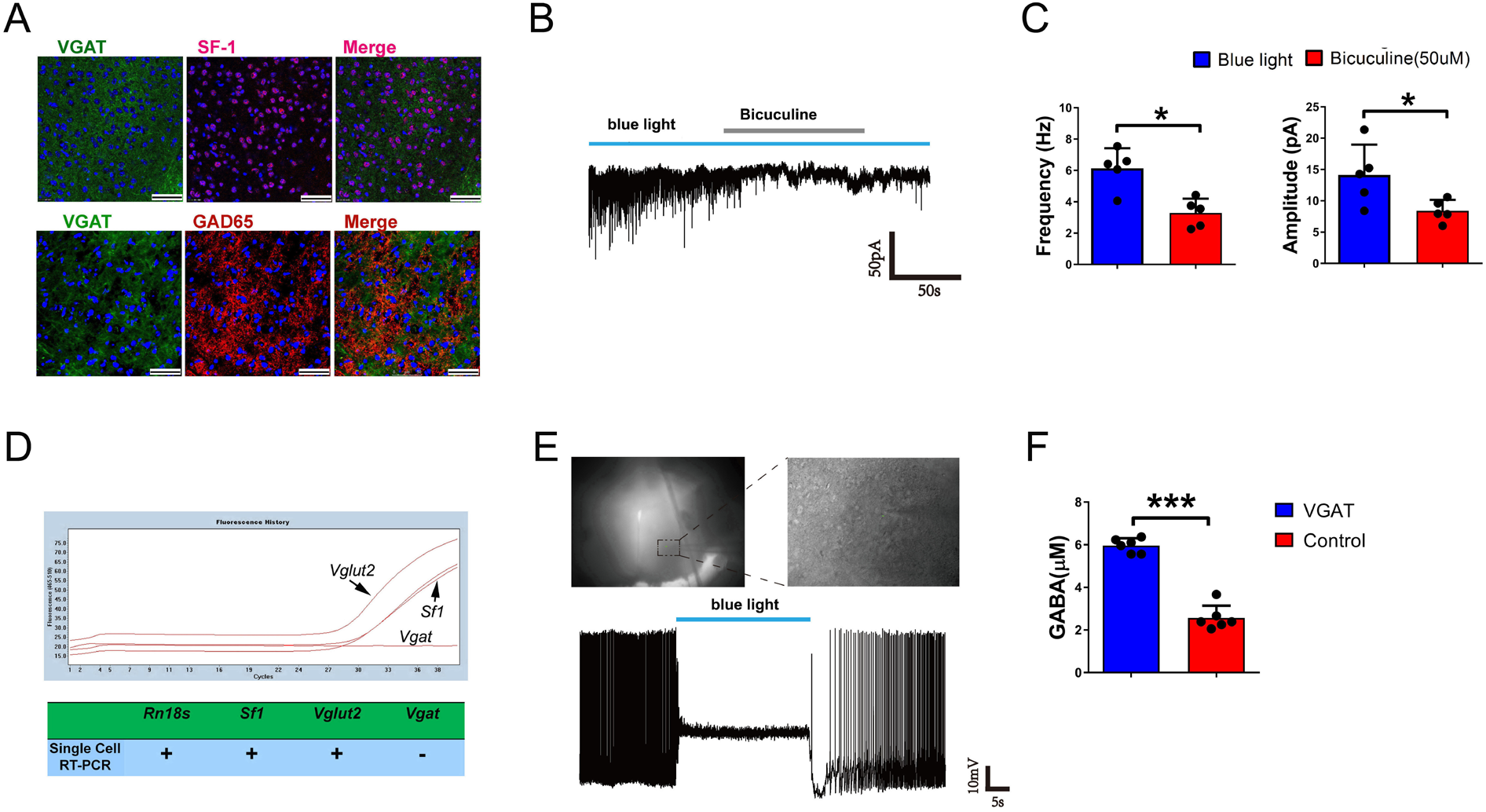
Activation of GABAergic projections in the VMHdm inhibit the firing of SF1 neurons. **(A)** High magnification image of VGAT signals surrounding SF1 neurons in the VMHdm region (top); double staining of VGAT and GAD65 revealed GABAergic axon terminals in VMHdm (bottom); scale bar, 50 μm. **(B)** An IPSC induced by blue light was blocked by bicuculine (50 μM). **(C)** Quantification of frequency and amplitude of IPSCs from SF1 neurons in the light stimulation and bicuculine groups; both the frequency and amplitude of the bicuculine group IPSCs was significantly lower; values represent mean±SEM (n=5 per group; \**p*<0.05; Student’s t test). **(D)** Single-cell RT-PCR of the patched cells revealed gene expression of *vglut2*, *Sf1* but not *vgat* (n=5). **(E)** Representative electrophysiological recording of the spontaneous firing of SF1 neurons after stimulating the VMHdm with blue light in VGAT mice. **(F)** Quantification of GABA levels in VMHdm of VGAT and control groups; values represent mean±SEM (n=6 per group; \*\*\**p*<0.001; Student’s t test).

**Fig. S3.**
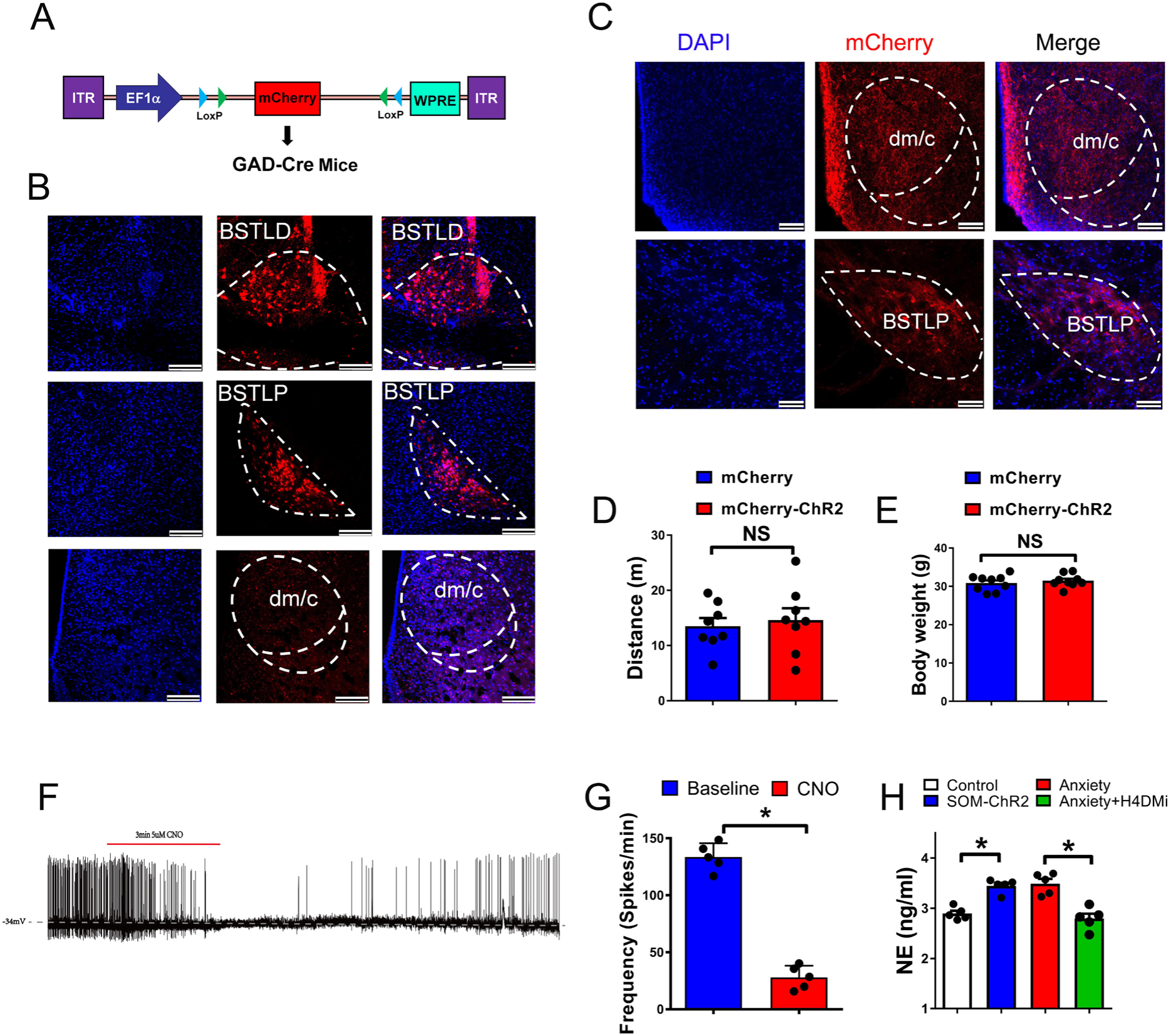
Somatostain neurons in the BNST send GABAergic projections to the VMHdm region. **(A)** Schematic showing AAV-Ef1a-DIO-mcherry virus injections into the lateral dorsal and lateral posterior region of BNST in GAD-Cre mice. **(B)** GABAergic neurons were observed in the BSTLD and the BSTLP; GABAergic neural projections were observed in the VMHdm (BSTLD, lateral-dorsal region of BNST; BSTLP, lateral posterior region of BNST); scale bar, 150 μm **(C)** Representative image of SOM positive neural projections in the VMHdm (top), and SOM positive neuron bodies in the BSTLP (bottom) (BSTLP, LP region of BNST); scale bar, 100 μm. **(D)** Total distance traveled in open field test showing mCherry-ChR2 and mCherry groups; values represent mean±SEM (n=8 per group; NS, not significant; Student’s t test). **(E)** Bodyweight of mice in the mCherry-ChR2 and mCherry groups; values represent mean±SEM (n=8 per group; NS, not significant; Student’s t test). **(F)** Representative electrophysiological recordings of somatostatin (SOM) neurons selectively silenced using DREADD technique. **(G)** Quantification of frequency (spikes/min) from SOM neurons in the baseline and CNO group. Values represent mean±SEM (n=5 per group; *p<0.05; Student’s t test). **(H)** Quantification of norepinephrine (NE) levels comparing SOM-ChR2, anxiety, anxiety + hM4Di and control groups; NE was significantly higher in both SOM-ChR2 and anxiety groups, however, it was significantly lower in hM4Di group; values represent mean±SEM (n=5 mice per group; \**p*<0.05; one-way analysis of variance (ANOVA) with Bonferroni correction for multiple comparisons test).

**Fig. S4.**
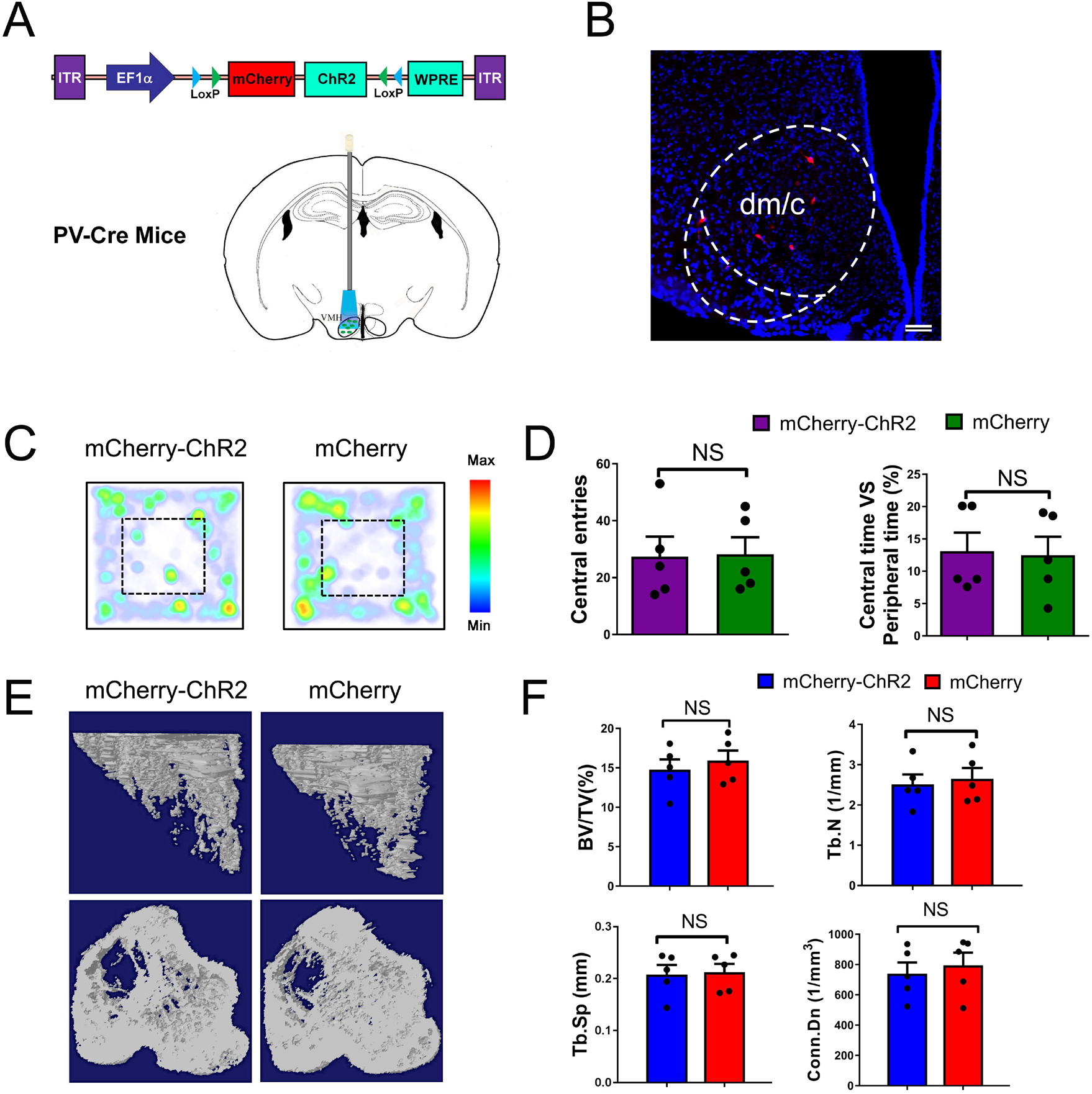
Activation of parvalbumin neural projections in the VMH did not induce anxiety-like behavior or bone loss. **(A)** Schematic showing AAV-Ef1α-DIO-ChR2-mcherry virus and parvalbumin-positive neural terminals were optogenetically activated in the VMHdm of PV-Cre mice. **(B)** The expression of ChR2-mcherry on parvalbumin neural projections; scale bar, 100 μm. **(C)** Open field test for the mCherry-ChR2 and mCherry groups. **(D)** Time in central area and central entries between the mCherry-ChR2 and mCherry groups; values represent mean±SEM (n=5 per group; NS, not significant; Student’s t test). **(E)** Representative micro-CT scans of bone structure in mCherry-ChR2 and mCherry groups. **(F)** MicroCT analysis of the parameters including trabecular bone volume/tissue volume (BV/TV), the trabecular number (TbN), trabecular separation (Tb.Sp), and connectivity density (Conn. Dn) in mCherry-ChR2 and mCherry groups; values represent mean± SEM (n=5 per group; NS, not significant; Student’s t test).

**Fig. S5.**
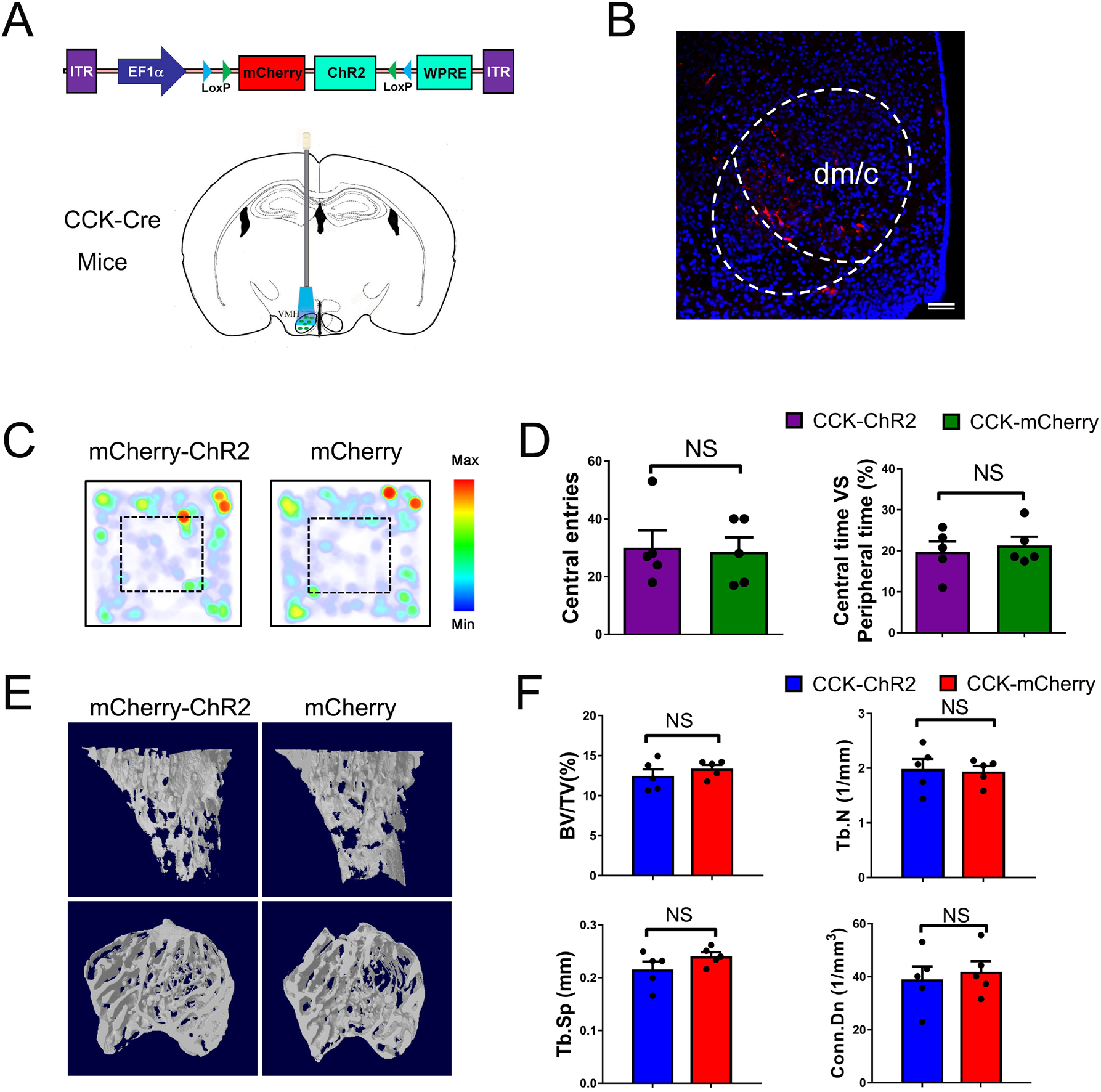
Activation of the Cholecystokinin neural projection in the VMH did not induce anxiety-like behavior or bone loss. **(A)** Schematic showing AAV-Ef1α-DIO-ChR2-mcherry and Cholecystokinin (CCK)-positive neural terminals were activated in the VMHdm of CCK-Cre mice. **(B)** Expression of ChR2-mcherry on cholecystokinin neural projections; scale bar, 100 μm. **(C)** Open field test comparing mCherry-ChR2 and mCherry groups. **(D)** Time spent by in the central area and central entries mCherry-ChR2 and mCherry groups; values represent mean±SEM (n=5 per group; NS, not significant; Student’s t test). **(E)** Representative micro-CT scans of bone structure in the mCherry-ChR2 and mCherry groups. **(F)** MicroCT analysis of the parameters including trabecular bone volume/tissue volume (BV/TV), the trabecular number (TbN), trabecular separation (Tb.Sp), and connectivity density (Conn. Dn) in the CCK-ChR2 and CCK-mCherry groups; values represent mean±SEM (n=5 per group; NS, not significant; Student’s t test).

**Fig. S6.**
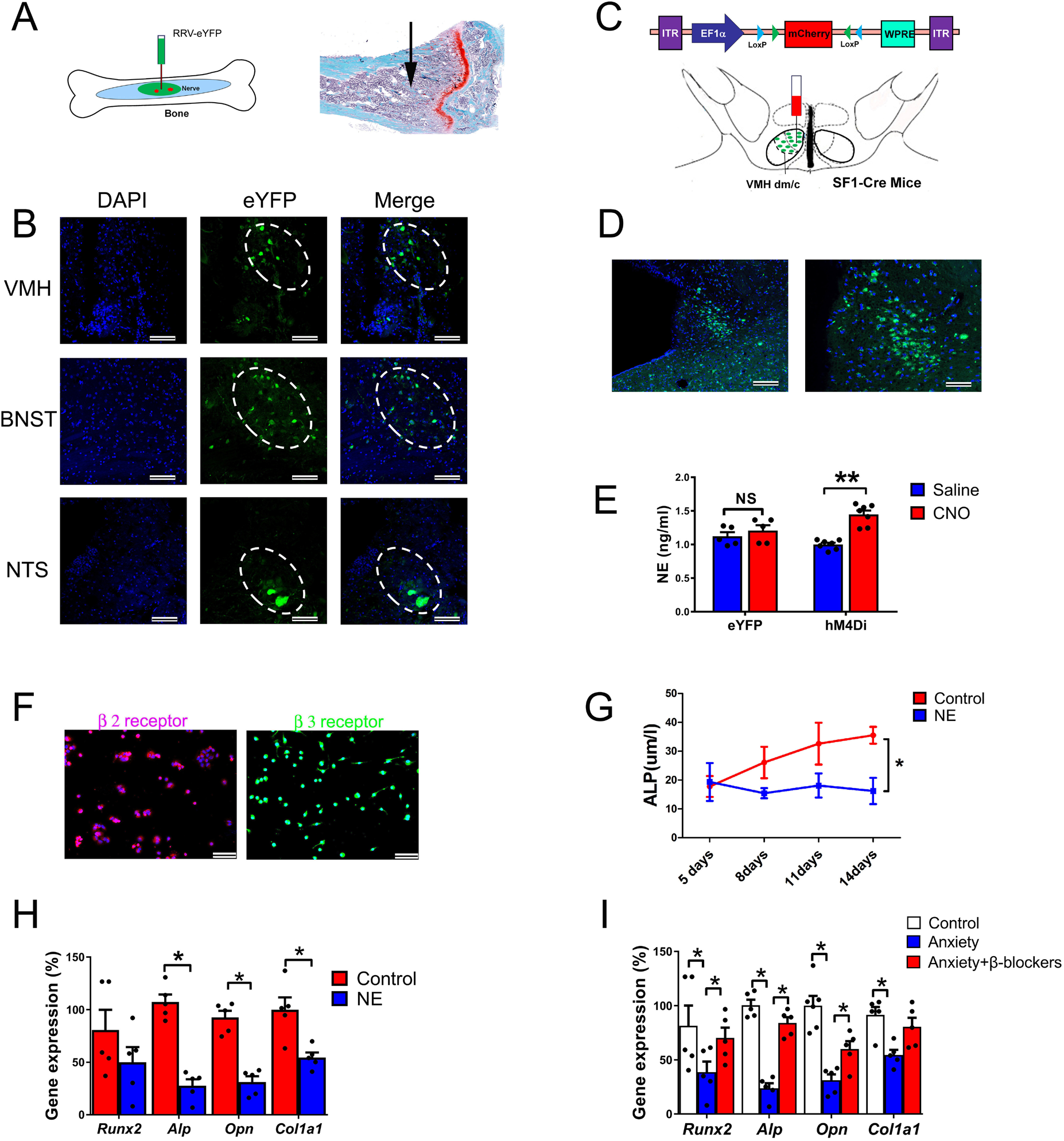
Sympathetic system and NE-mediated BNST-VMH-NTS neural circuitry-induced bone loss. **(A)** Schematic showing poly-synaptic retrograde PRV virus injected into murine bone marrow to label the neurons innervating the bone (left); safranin-O staining of the mouse tibia (right). **(B)** eYFP labeled neurons were observed in the ventromedial hypothalamus (VMH), bed nucleus of the stria terminalis (BNST) and nucleus of the solitary tract (NTS); scale bar, 50 μm. **(C)** Schematic showing AAV expressing mCherry under the EF1α promoter; virus was injected into the dorsomedial region of the VMH in SF-Cre mice. **(D)** Expression of hM4di (green) in Vglut2 neurons in solM region of NTS; scale bar, 100 μm (left); 50 μm (right). **(E)** Quantification of norepinephrine (NE) levels in the eYFP and hM4Di groups; NE was significantly higher in hM4Di group compared to the saline group after CNO injection; values represent mean±SEM (n=5 mice per group; NS, not significant; \*\**p*<0.01; one-way analysis of variance (ANOVA) with Bonferroni correction for multiple comparisons test). **(F)** Immunostaining of β2 and β3 adrenergic receptors on osteoprogenitor cells; scale bar, 50 μm. **(G)** Quantification of alkaline phosphatase (ALP) during osteoblastic differentiation of osteoprogenitors in NE and control groups; values represent mean±SEM (n=5 per group; NS, not significant; Student’s t test). **(H)** RT-PCR analysis of *Runx2*, *Alp*, *Col1a1*, and *Opn* in the NE and control groups after 2 weeks of the osteogenic differentiation of osteoprogenitor cells; values represent mean±SEM (n=5 mice per group; \**p*<0.05; one-way analysis of variance (ANOVA) with Bonferroni correction for multiple comparisons test). **(I)** Gene expression analysis of *Runx2*, *Alp*, *Col1a1*, and *Opn* using freshly isolated bone marrow cells from mice in control group, anxiety group and anxiety + β-blockers group; values represent mean±SEM (n=5 mice per group; \**p*<0.05; one-way analysis of variance (ANOVA) with Bonferroni correction for multiple comparisons test).

**Fig. S7.**
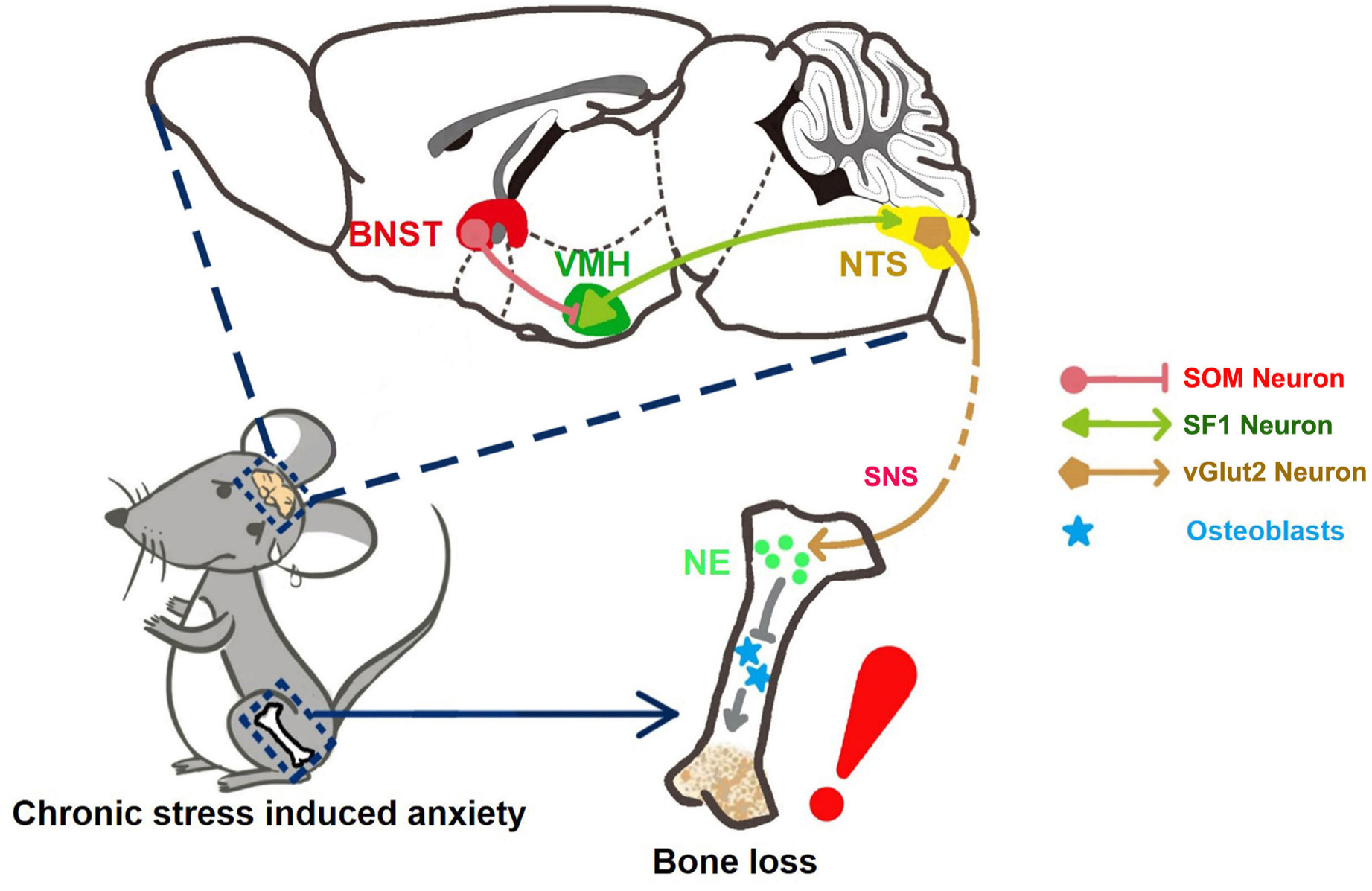
Schematic showing that the neural circuit BNST-VMHNTS can regulate anxiety-induced bone loss

